# The Role of the Tyrosine-Based Sorting Signals of the ORF3a Protein of SARS-CoV-2 on Intracellular Trafficking, Autophagy, and Apoptosis

**DOI:** 10.1101/2023.07.24.550379

**Authors:** Wyatt Henke, Maria Kalamvoki, Edward B Stephens

## Abstract

The open reading frame 3a (ORF3a) is an accessory transmembrane protein that is important to the pathogenicity of SARS-CoV-2. The cytoplasmic domain of ORF3a has three canonical tyrosine-based sorting signals (YxxΦ; where x is any amino acid and Φ is a hydrophobic amino acid with a bulky -R group). They have been implicated in the trafficking of membrane proteins to the cell plasma membrane and to intracellular organelles. Previous studies have indicated that mutation of the ^160^YSNV^163^ motif abrogated plasma membrane expression and inhibited ORF3a-induced apoptosis. However, two additional canonical tyrosine-based sorting motifs (^211^YYQL^213^, ^233^YNKI^236^) exist in the cytoplasmic domain of ORF3a that have not been assessed. We removed all three potential tyrosine-based motifs and systematically restored them to assess the importance of each motif or combination of motifs that restored efficient trafficking to the cell surface and lysosomes. Our results indicate that the YxxΦ motif at position 160 was insufficient for the trafficking of ORF3a to the cell surface. Our studies also showed that ORF3a proteins with an intact YxxΦ at position 211 or at 160 and 211 were most important. We found that ORF3a cell surface expression correlated with the co-localization of ORF3a with LAMP-1 near the cell surface. These results suggest that YxxΦ motifs within the cytoplasmic domain may act cooperatively in ORF3a transport to the plasma membrane and endocytosis to lysosomes. Further, our results indicate that certain tyrosine mutants failed to activate caspase 3 and did not correlate with autophagy functions associated with this protein.

**IMPORTANCE:** Open reading frame 3a (ORF3a) encodes for the largest of the SARS-CoV-2 accessory proteins. While deletion of the ORF3a gene from SARS-CoV-2 results in a virus that replicates slightly less efficiently in cell culture, deletion also results in a virus that is less pathogenic in mouse models of SARS-CoV-2 infections. The ORF3a has been reported to be a viroporin, induces apoptosis and incomplete autophagy in cells. Thus, determining the domains involved in these functions will further our understanding of how this protein influences virus assembly and pathogenesis. Here, we investigated the role of the three potential tyrosine-based sorting signals in the cytoplasmic domain of the ORF3a on intracellular protein trafficking, apoptosis, and in the initiation of autophagy. Our results indicate that more than one YxxΦ motif is required for efficient transport of ORF3a, ORF3a expression resulted in minimal apoptosis, and cell surface expression was not required for autophagy.

## INTRODUCTION

First isolated from SARS-CoV, the ORF3a protein is the largest of the SARS-CoV-2 accessory proteins having a N-terminal domain of approximately 40 amino acids, three transmembrane domains, and a longer 150 amino acid cytoplasmic domain (**1**). The ORF3a proteins from SARS-CoV and SARS-CoV-2 form multimeric structures in membranes that have been reported to be ion channels (i.e., viroporins) that was previously shown to be permissive to divalent cations (**2-4**). The SARS-CoV-2 ORF3a behaved as a cation channel with a large single-channel conductance (375 pA) and had a modest selectivity for Ca^++^ and K^+^ over Na^+^. However, the SARS-CoV-2 channel was not blocked by Ba^++^ as was the case for the SARS-CoV channel (**2**). A more recent study indicates that ORF3a may not actually be a viroporin (**5**). ORF3a also has been shown to induce apoptosis and incomplete autophagy (**6,7**). Ectopic expression of ORF3a revealed that it is readily expressed at the cell plasma membrane and in several intracellular compartments of the cell including the ER, ERGIC, Golgi complex, *trans* Golgi network (TGN), and lysosomes.

It has been established that linear sorting signals mediate trafficking of cellular membrane proteins from the Golgi complex to their final membrane compartment. Most of this sorting occurs in the TGN (**8**). These short linear sequences include the: a) dileucine motifs ([D/E]xxxL[L/I] or [D/E]xxL[L/I] ; b) tyrosine-based motifs (YxxΦ; with x being any amino acid and Φ being an amino acid with large hydrophobic -R group) and are both recognized by adaptor protein complexes AP-1, AP-2, and AP-3; and c) Asn-Pro-X-Tyr (NPXY) motifs, which are recognized by the accessory clathrin adaptor proteins (**9, 10**). The YxxΦ sorting signals have dual specificity directing the trafficking of membrane proteins within the endosomal and/or secretory pathways and also in rapid endocytosis from the cell surface and/or sorting to lysosomes and lysosome-related organelles (**11-18**).

Like cellular membrane proteins, many viral membrane proteins also use canonical tyrosine-based signals for intracellular transport and mutation of these sites can affect trafficking of these proteins to sites of assembly (**9**). While the spike (S) and envelope (E) proteins of SARS-CoV-2 have no YxxΦ motifs, both the membrane (M) protein and viroporin ORF3a have YxxΦ motifs. Further, previous studies on the ORF3a proteins of SARS-CoV and SARS-CoV-2, indicate that the mutation of the tyrosine residue in motif ^160^YNSV^163^ resulted in an ORF3a that was transported to the Golgi complex but was not present at the cell surface (**6, 19**). Further, these investigators showed that the lack of cell surface expression correlated with a reduction in apoptosis (**6, 19**). However, the cytoplasmic domains from ORF3a proteins from the SARS-CoV, SARS-CoV-2, and various related bat coronaviruses within the genus Sarbecovirus of the β-coronaviruses have two to three well-conserved YxxΦ motifs in the cytoplasmic domain. Here, we have performed a detailed analysis of the intracellular trafficking and potential role in apoptosis of the three potential YxxΦ sorting motifs of the SARS-CoV-2 ORF3a. Our results indicated that an ORF3a with a single tyrosine motif ^160^YNSV^163^ was insufficient for trafficking to the plasma membrane and induction of apoptosis. Our findings indicate that the YYQL motif at position 211-214 was most important in targeting ORF3a to the cell plasma membrane. Further, our studies on ORF3a and the tyrosine-based sorting signal mutants indicate that removal of all tyrosine-based motifs inhibited procaspase 3 cleavage and LC3 lipidation but not p62 degradation.

## RESULTS

### The ORF3a has more than one potential tyrosine-based sorting motif

We analyzed the ORF3a sequences from SARS-CoV-2 and SARS-CoV-2-like viruses (275 amino acids in length) for tyrosine-based sorting motifs (YxxΦ). The results indicated that at least three potential tyrosine-based sorting motifs were found in the cytoplasmic domain (**Fig. 1**). All isolates had a conserved ^160^**YSNV**^163^ motif but also had two additional motifs at (^211^**YYQL**^213^) and (^233^**YNKI**^236^). We also observed that the ORF3a proteins from SARS-CoV and SARS-CoV-like viruses (274 amino acids in length) also had two to three motifs (**Supplemental Fig. 1**).

**Figure 1.**
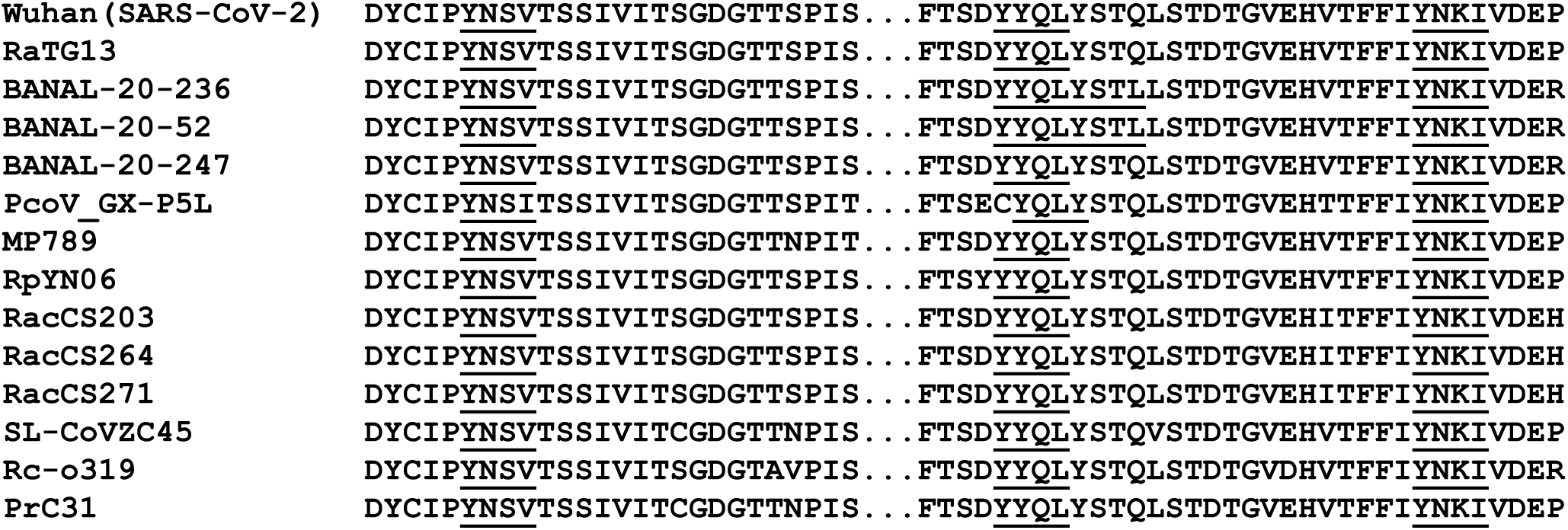
Alignment of SARS-CoV-2 and SARS-CoV-2-like ORF3a sequences (amino acids 155-240) with potential tyrosine-based sorting signals (YxxΦ) in the cytoplasmic domains underlined and bolded. The isolate, (species), and NIH accession numbers are: 1) Wuhan strain (Homo sapiens; accession #P0DTC3); 2) RATG13 (*Rhinolophus affinis*; accession #QHR63301); 3) Banal-20-236 (*Rhinolophus marshalli* ;accession # UAY13254); 4) Banal 20-52 (*Rhinolophus marshalli*; accession #UAY13218; 5) Banal20-247 (*Rhinolophus malayanus*; accession #UAY13266); 6) PcoV_GX-P5L (*Manis javanica*; accession # QIA48633); 6) MP789 (*Manis javanica*) accession #QIG55946; 7) MP796 (*Manis javanica*, accession #QIG55946; 8) RpYN06 (*Rhinolophus pusillus;* accession # QWN56253); 9) RacCS203 (*Rhinolophus acuminatus*; accession #QQM18865); 10) RacCS264 (*Rhinolophus acuminatus*; accession #QQM18898); 11) RacCS271 (*Rhinolophus acuminatus*; accession #QQM18909); 12) SL-CoVZC45 (*Rhinolophus pusillus*; accession #AVP78032); 13) Rc-o319 (*Rhinolophus cornutus*; accession #BCG66628); and 14) PrC31 (*Rhinolophus blythi*; accession #QSQ01651). The potential tyrosine-based sorting signals are bolded and underlined.

### Expression of SARS-CoV-2 ORF3a and various mutants

We generated a series of ORF3a mutants in which one, two, or all three tyrosine residues of these potential sorting signals were changed to alanine residues. These mutants included one in which all three tyrosine were altered to alanine (ORF3a-ΔYxxΦ), those mutants in which two tyrosine residues were substituted with alanines (ORF3a-Y160, ORF3a-Y211, and ORF3a-Y233), and those in which one tyrosine residue was substituted with alanines (ORF3a-Y160,211, ORF3a-Y160,233, and ORF3a-Y211,233). For the designation of the six mutants above, the number following the Y indicates the tyrosine motif(s) that were intact. The unmodified ORF3a and mutants all had an HA-tag at the N-terminus (**Fig. 2A**). We examined the steady-state levels to determine if the different mutations in ORF3a protein were stably expressed in cells or if were rapidly degraded. Vectors expressing the SARS-CoV-2 ORF3a or ORF3a mutants were transfected into HEK293 cells. Cell lysates were prepared at 48 h post-transfection and ORF3a proteins analyzed on immunoblots using an anti-HA antibody. Our results show that all eight ORF3a proteins were expressed well in comparable numbers of HEK293 cells (**Fig. 2B**).

**Figure 2.**
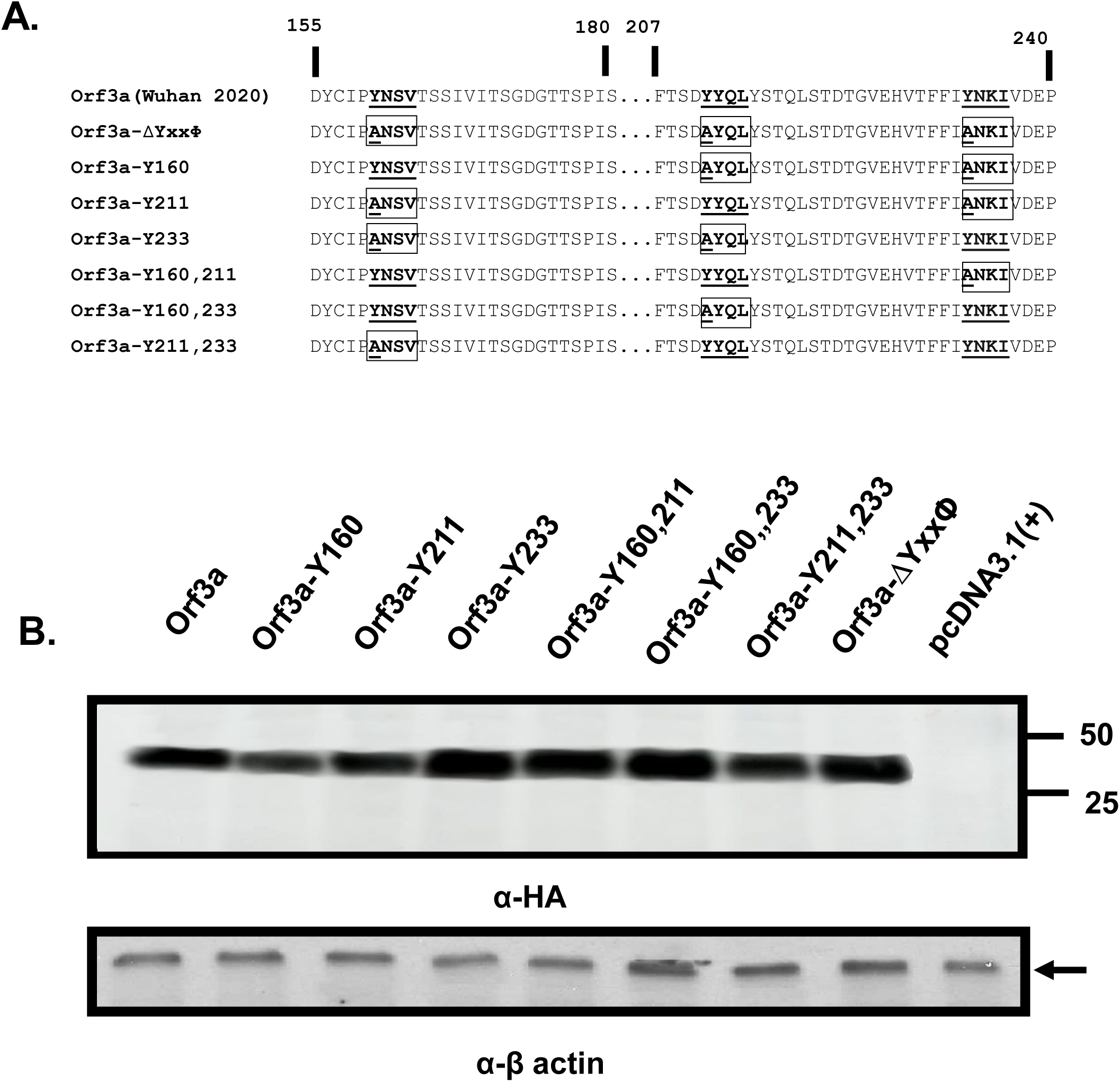
The ORF3a mutants analyzed in this study. **Panel A**. The unmodified ORF3a and the mutants were constructed for this study. The potential tyrosine-based sorting signals that were unchanged were bolded and underlined. Those potential tyrosine-based sorting signals that were altered are boxed and bolded with the amino acid changes underlined. **Panel B**. Expression of the ORF3a and its mutants. HEK293 cells were transfected with vectors expressing the unmodified ORF3a and the seven mutants were analyzed. Proteins were separated by SDS-PAGE, transferred to membranes, and analyzed in immunoblots using an antibody directed against the C-terminal HA-tag. β- actin served as a control for loading of samples.

### The unmodified ORF3a protein is transported through the secretory pathway to cell plasma membrane

To determine if ORF3a proteins were transported to the ER, cis medial Golgi, the *trans* Golgi network (TGN), or the mitochondria, cells were co-transfected with the vector expressing the ORF3a protein and vectors expressing each of the intracellular markers as described in the Materials and Methods section. With these co-transfections, cells were fixed at 48 h post-transfection and immunostained for ORF3a using an anti-HA antibody followed by a reaction with an appropriate secondary antibody as described in the Materials and Methods section. Our results indicate that the unmodified ORF3a co-localized with markers for the ER, ERGIC, and TGN (**Fig. 3**) but not with the 4xmts-mNeonGreen mitochondrial marker (data not shown). Our also showed that the ORF3a also co-localized with markers for the cis-medial Golgi (Giantin) and trans Golgi (Golgin 97).

**Figure 3.**
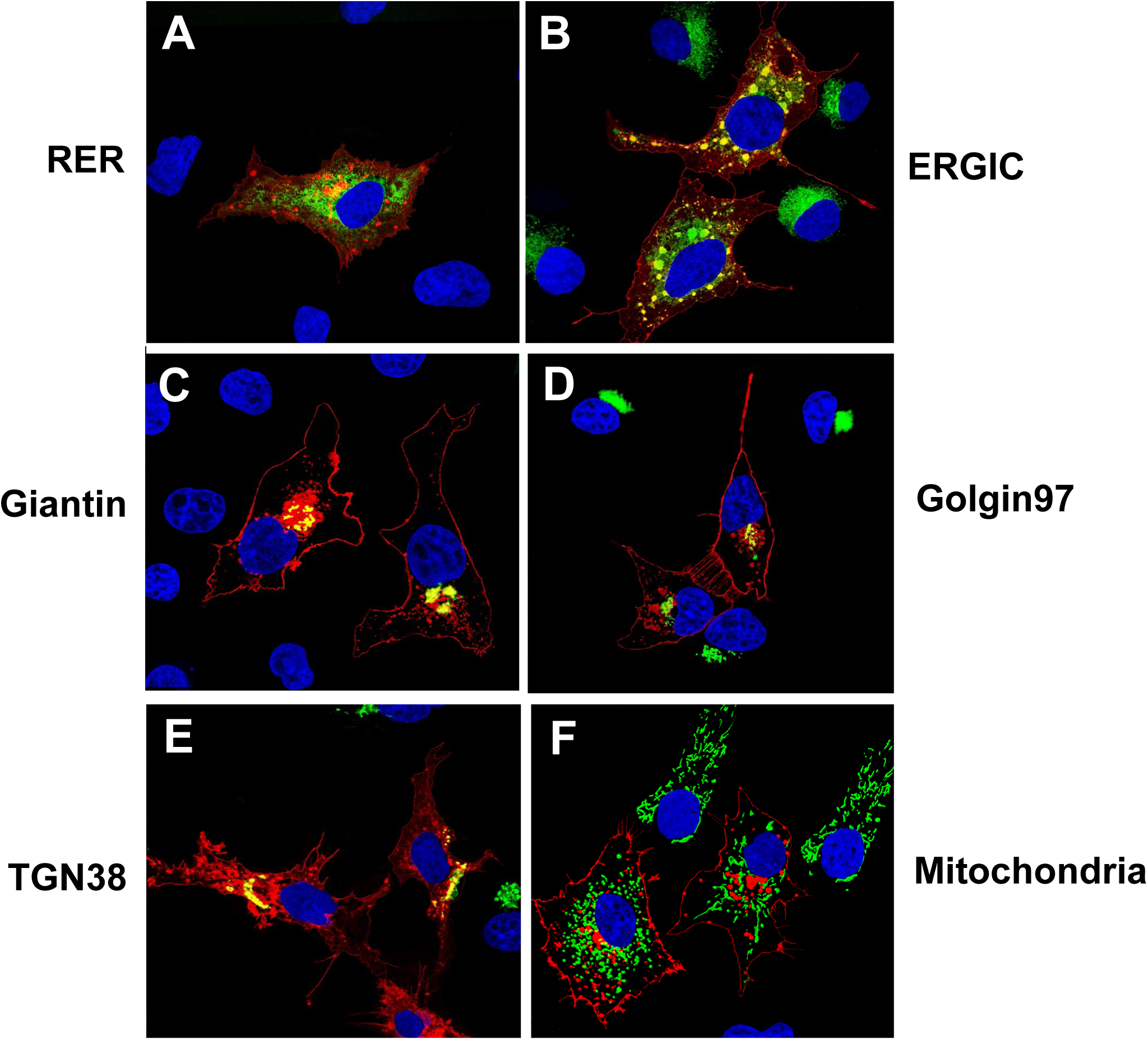
The SARS-CoV-2 ORF3a is expressed in organelles of the secretory pathway and at the cell plasma membrane. COS-7 cells were co-transfected with the empty pcDNA.3.1(+) vector or vectors expressing SARS-CoV-2 ORF3a-HA protein and vectors expressing markers for the rough endoplasmic reticulum (ER-MoxGFP), and *trans* Golgi network (TGN38GFP) and mitochondria (4xmts-mNeonGreen. In other cultures, COS-7 cells were transfected with the pcDNA.3.1(+) vectors expressing SARS-CoV-2 ORF3a-HA and immunostained with antibodies against other intracellular organelles (ERGIC or Golgin 97) as described in the Materials and Methods. At 48 h post transfection, cells were fixed, permeabilized, and blocked. The coverslips were reacted with a mouse monoclonal antibody against the HA-tag of HA-ORF3a and with a rabbit antibody against ERGIC53 (ERGIC) or Golgin 97 (*trans* Golgi) followed by appropriate secondary antibodies, as described in the Materials and Methods section. Coverslips with cells were washed, counter stained with DAPI (1 μg/ml) and mounted. Cells were examined using a Leica TCS SPE confocal microscope. using a 100x objective with a 2x digital zoom using the Leica Application Suite X (LAS X, LASX) software package. A minimum of 100 cells were examined per staining with the micrographs shown being representative. Panel A. Cells transfected with vectors expressing ORF3a and ER-moxGFP. Panel B. Cells transfected with a vector expressing ORF3a and immunostained with antibodies against ERGIC-53 and HA. Panel C. Cells transfected with vectors expressing ORF3a and mNeonGreen-Giantin Panel D. Cells transfected with the vector expressing ORF3a and immunostained with antibodies against Golgin 97. Panel E. Cells transfected with vectors expressing ORF3a and TGN-38GFP. Panel F. Cells transfected with the vector expressing ORF3a and 4xmts-mNeon Green.

### The ORF3a-ΔYxxΦ mutant is poorly expressed at the cell surface

We next analyzed ORF3a-ΔYxxΦ for its intracellular localization with appropriate intracellular markers as we had been done for the unmodified ORF3a. We observed that this protein co-localized the ER, ERGIC, and TGN but was neither detected at the cell plasma membrane (**Fig. 4**) nor was it associated with mitochondria (data not shown). Our results also showed that the ORF3a-YxxΦ also co-localized with markers for the *cis*-*medial* Golgi and *trans* Golgi.

**Figure 4.**
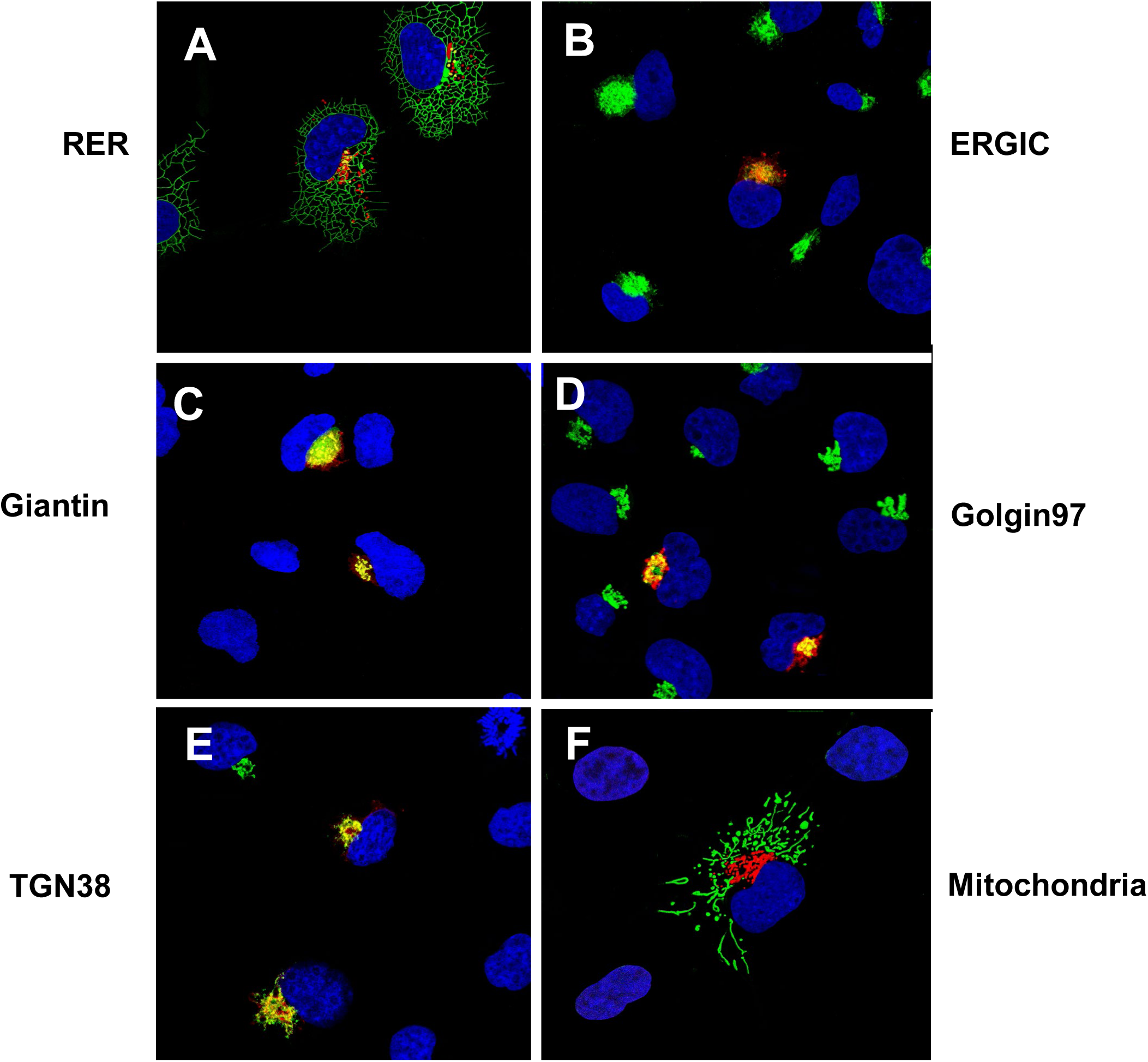
The ORF3a-ΔYxxΦ is not expressed at the cell plasma membrane. HEK293 cells were transfected with the empty pcDNA.3.1(+) vector or a vector expressing the SARS-CoV-2 HA-ORF3a-ΔYxxΦ protein section as in Figure 3. Panel A. Cells transfected with vectors expressing HA-ORF3a-ΔYxxΦ and ER-MoxGFP and immunostained with an anti-HA antibody. Panel B. Cells transfected with a vector expressing HA-ORF3a-ΔYxxΦ and immunostained with antibodies against ERGIC-53 and HA. Panel C. Cells transfected with vectors expressing HA-ORF3a-ΔYxxΦ and mNeonGreen-Giantin and immunostained with an anti-HA antibody. Panel D. Cells transfected with the vector expressing HA-ORF3a-ΔYxxΦ and immunostained with antibodies against Golgin 97 and HA. Panel E. Cells transfected with vectors expressing HA-ORF3a-ΔYxxΦ and TGN-38GFP and immunostained with an anti-HA antibody. Panel F. Cells transfected with the vector expressing HA-ORF3a-ΔYxxΦ and 4xmts-mNeonGreen and immunostained with antibodies against HA.

### Intracellular expression of ORF3a mutants with one potential tyrosine-based sorting motifs intact

We next examined the intracellular localization of ORF3a mutants with one potential intact tyrosine-based motif (ORF3a-Y160, ORF3a-Y211, and ORF3a-Y233). (**Fig. 5A-I**). Co-transfection of vectors expressing the ORF3a-Y160 (Panels A-C) or ORF3a-Y233 (Panels G-I) with vectors expressing either ER-moxGFP or TGN38-EGFP fusion proteins revealed that both proteins co-localized with the ER and TGN markers but were not observed at the cell plasma membrane (**Fig. 5A-C, G-I**). Transfection of cells with the vector expressing ORF3a-Y211 revealed that this protein co-localized with the ER and TGN markers and was readily detectable at the cell surface (**Fig. 5D-F**).

**Figure 5.**
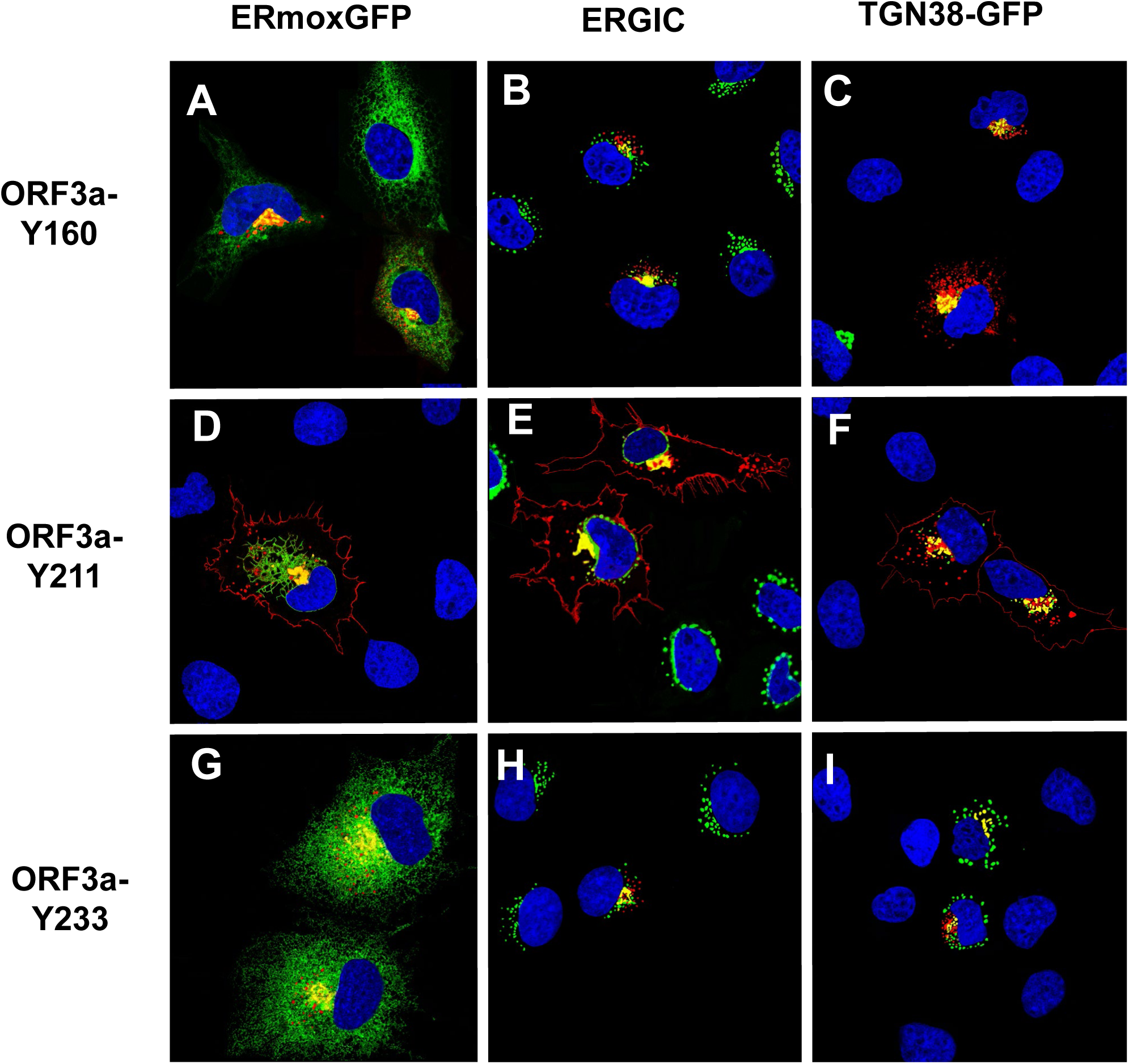
Cell expression of ORF3a mutants with one intact tyrosine-based motif intact. COS-7 cells were co-transfected with vectors expressing HA-ORF3a, HA-ORF3a-Y160, HA-ORF3a-Y211, or HA-ORF3a-Y233 and a vector expressing ER-moxEGFP or TGN38-EGFP. At 48 h post-transfection, cells were fixed, permeabilized, and blocked. Cells were reacted with a mouse monoclonal antibody against the HA-tag and overnight, washed, and reacted with an appropriate secondary antibody tagged with Alexa Fluor 594 (for HA) for 1 h. Cells were washed and counter-stained with DAPI (1 μg/ml) for 5 min. Cells were viewed using a Leica TC8 confocal microscope as described in the Materials and Methods section. At least 50 cells were examined for surface expression and co-localization with ERmoxEGFP or TGN38-EGFP. Panel A. Cells transfected with a vector expressing HA-ORF3a-Y160 and ERmoxEGFP and immunostained with antibodies against the HA-tag. Panel B. Cells transfected with a vector expressing HA-ORF3a-Y160 and immunostained with antibodies against the HA-tag and ERGIC-53. Panel C. Cells transfected with a vector expressing HA-ORF3a-Y160 and TGN38-EGFP and immunostained with an antibody against the HA-tag. Panel D. Cells transfected with vectors expressing HA-ORF3a-Y211 and ERmoxEGFP and immunostained with antibodies against the HA-tag. Panel E. Cells transfected with a vector expressing HA-ORF3a-Y160 and immunostained with antibodies against the HA-tag and ERGIC-53. Panel F. Cells transfected with a vector expressing HA-ORF3a-Y211 and TGN38-EGFP and immunostained with antibodies against the HA-tag. Panel G. Cells transfected with vectors expressing HA-ORF3a-Y233 and ERmoxEGFP and immunostained with antibodies against the HA-tag. Panel H. Cells transfected with vectors expressing HA-ORF3a-Y233 and immunostained with antibodies against the HA-tag and ERGIC-53. Panel I. Cells transfected with a vector expressing HA-ORF3a-Y160 and TGN-EGFP and immunostained with antibodies against the HA-tag.

### Intracellular expression of ORF3a mutants with two potential tyrosine-based sorting motifs intact

We next analyzed the ORF3a mutants with two intact tyrosine motifs (ORF3a-Y160,211; ORF3a-Y160,233; and ORF3a-Y211,233). The rationale behind analyzing these mutants was to determine if more than one tyrosine motif was required for efficient trafficking to the cell surface. Our results indicated that of these three mutants, only ORF3a-Y160,211 was observed at the cell surface while ORF3a-Y160,233 and ORF3a-Y211,233 were associated with intracellular compartments (**Fig. 6A-I**).

**Figure 6.**
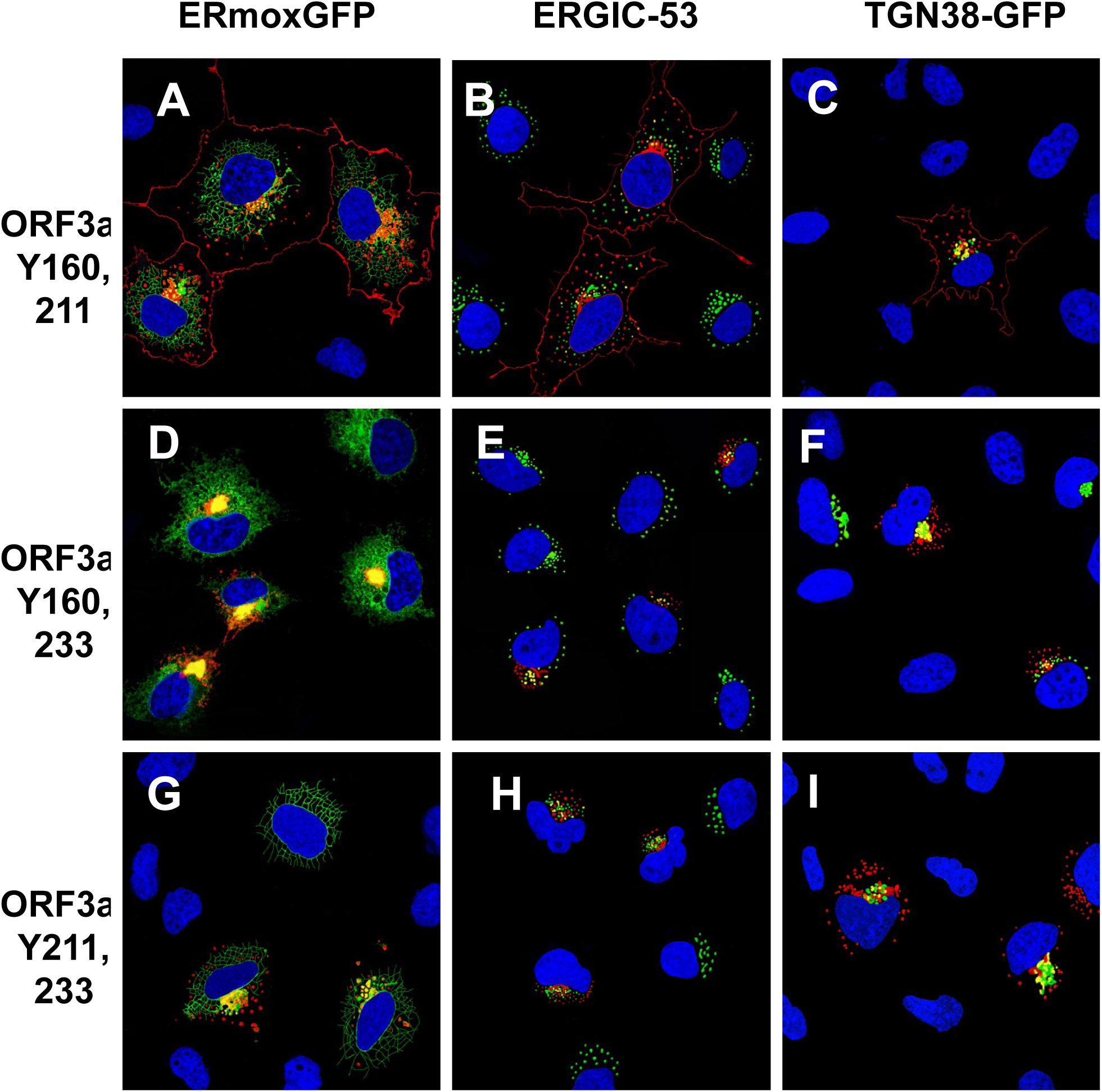
Cell expression of ORF3a mutants with two intact tyrosine motifs intact. COS-7 cells were co-transfected with vectors expressing HA-ORF3a, HA-ORF3a-Y160,211, HA-ORF3a-Y160,233, or HA-ORF3a-Y211,233. At 48 h post-transfection, cells were fixed, permeabilized, and blocked. Cells were reacted with a mouse monoclonal antibody against the HA-tag overnight, washed, and reacted with an appropriate secondary antibody tagged with Alexa Fluor 594 for HA for 1 h. Cells were washed and counter-stained with DAPI (1 μg/ml) for 5 min. Cells were viewed using a Leica TC8 confocal microscope as described in the Materials and Methods section. At least 50 cells were examined for expression and co-localization. **Panel A.** Cells transfected with a vector expressing HA-ORF3a-Y160,211 and ERmoxEGFP and immunostained with antibodies against the HA-tag. **Panel B.** Cells transfected with a vector expressing HA-ORF3a-Y160,211 and immunostained with antibodies against the HA-tag and ERGIC-53. **Panel C**. Cells transfected with a vector expressing HA-ORF3a-Y160,211 and TGN38-EGFP and immunostained with antibodies against the HA-tag. **Panel D.** Cells transfected with vectors expressing HA-ORF3a-Y160,233 and ERmoxEGFP and immunostained with antibodies against the HA-tag. **Panel E**. Cells transfected with a vector expressing HA-ORF3a-Y160,233 and immunostained with antibodies against the HA-tag and ERGIC-53. **Panel F**. Cells transfected with a vector expressing HA-ORF3a-Y160,211 and TGN38-EGFP and immunostained with antibodies against the HA-tag. **Panel G**. Cells transfected with vectors expressing HA-ORF3a-Y211,233 and ERmoxEGFP and immunostained with antibodies against the HA-tag. **Panel H**. Cells transfected with a vector expressing HA-ORF3a-Y211,233 and immunostained with antibodies against the HA-tag and ERGIC-53. **Panel H**. Cells transfected with vectors expressing HA-ORF3a-Y211,233 and TGN38-EGFP and immunostained with antibodies against the HA-tag.

#### Surface immunostaining of cells transfected with vectors expressing the ORF3a and ORF3a mutants confirm the cell surface expression patterns

To confirm that ORF3a, ORF3a-Y211, and ORF3a-160,211 were expressed at the cell surface, we also performed a double immunostaining assay, relying on sequential treatment with antibodies before and after cell permeabilization, as described in the Materials and Methods. Both immunostainings were performed on ice. If the ORF3a or various mutants were transported to the cell surface, the cells should stain with the secondary Alexa Fluor 594 while internal ORF3a proteins should stain with the secondary antibody tagged with AlexaFluor488. Conversely, if the ORF3a protein was not expressed on the cell surface, immunostaining with the anti-HA and the Alexa Fluor594 should yield little to no red staining while the cells should stain internally with anti-HA and the secondary antibody tagged with Alexa Fluor488. All micrographs were taken using the same exposure time and laser intensity. As we expected, cells transfected with vectors expressing the unmodified HA-ORF3a were stained at the cell surface (evidenced by the red color) while cells transfected with the vector expressing HA-ORF3a-YxxΦ had virtually no immunostaining at the cell surface (**Fig. 7B-D**). Both constructs exhibited internal staining (**Fig. 7D-E**) and merging images taken at 488nm and 594nm confirmed the surface staining of HA-ORF3a but not HA-ORF3a-YxxΦ (**Fig. 7C, F**). Using the same methodology, mutants HA-ORF3a-Y160, HA-ORF3a-Y233, HA-ORF3a-Y160,233, and HA-ORF3a-Y211,233 were not observed on the cell surface while both HA-ORF3a-Y211 and HA-ORF3a-Y160,211 were detectable at the cell surface (**Fig. 8-9**). Taken together, these results indicate that: a) the tyrosine motif ^160^YNSV^163^ alone is insufficient for the transport of ORF3a to the cell surface; b) the tyrosine motif ^211^YYQL^214^ alone can target ORF3a to the cell surface; c) the ORF3a with the tyrosine motifs ^160^YNSV^163^ and ^211^YYQL^214^ were transported to the cell plasma membrane; and d) the presence of the tyrosine motif ^233^YNKI^236^ with a tyrosine motif at positions 160-163 or 211-214 was not efficiently transported to the cell surface.

**Figure 7.**
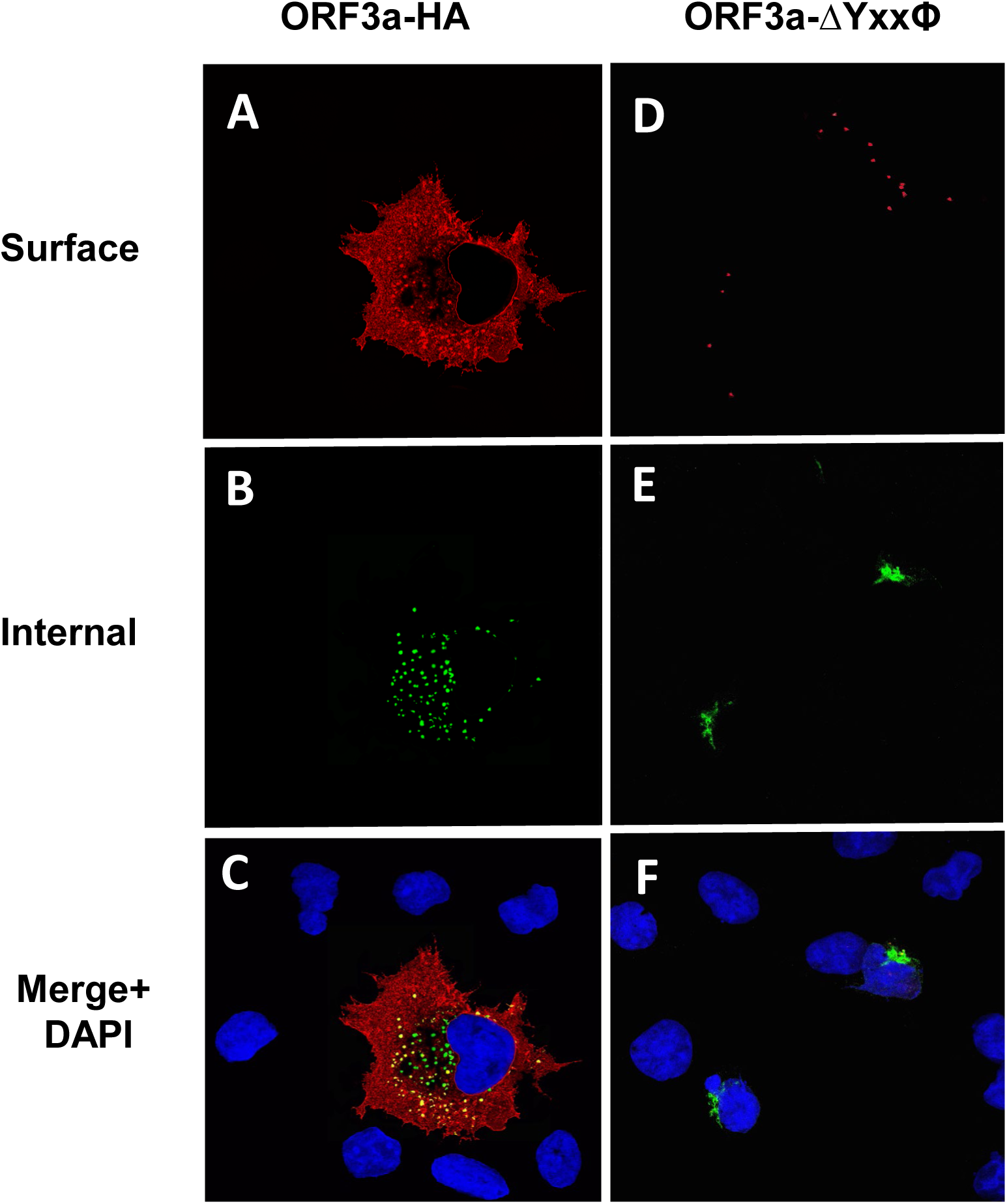
Surface immunostaining of cells transfected with vectors expressing ORF3a and ORF3a-YxxP. COS-7 cells were transfected with vectors expressing either ORF3a (panels A-C) or ORF3a-YxxP (Panels D-F). COS-7 cells transfected with the empty vector showed no immunofluorescence (data not shown). At 24 h post-transfection, cells were immunostained with an antibody against the HA-tag followed by a secondary antibody tagged with Alexa Fluor 594. Cells were washed three times and permeabilized as described in the Materials and Methods section. Permeabilized cells were then reacted with the same primary antibody and a secondary antibody tagged with Alexa Fluor 488. The cells were washed, mounted, and examined using a Leica TSP8 laser scanning confocal microscope. Panel A. Surface immunostaining of ORF3a with an antibody against the HA-tag. Panel B. Internal immunostaining of ORF3a with an antibody against the HA-tag. Panel C. Merge of Panels A and B. Panel D. Surface immunostaining of ORF3a-YxxΦ with an antibody against the HA-tag. Panel E. Internal immunostaining of ORF3a-YxxΦ with an antibody against the HA-tag. Panel F. Merge of Panels D and E.

**Figure 8.**
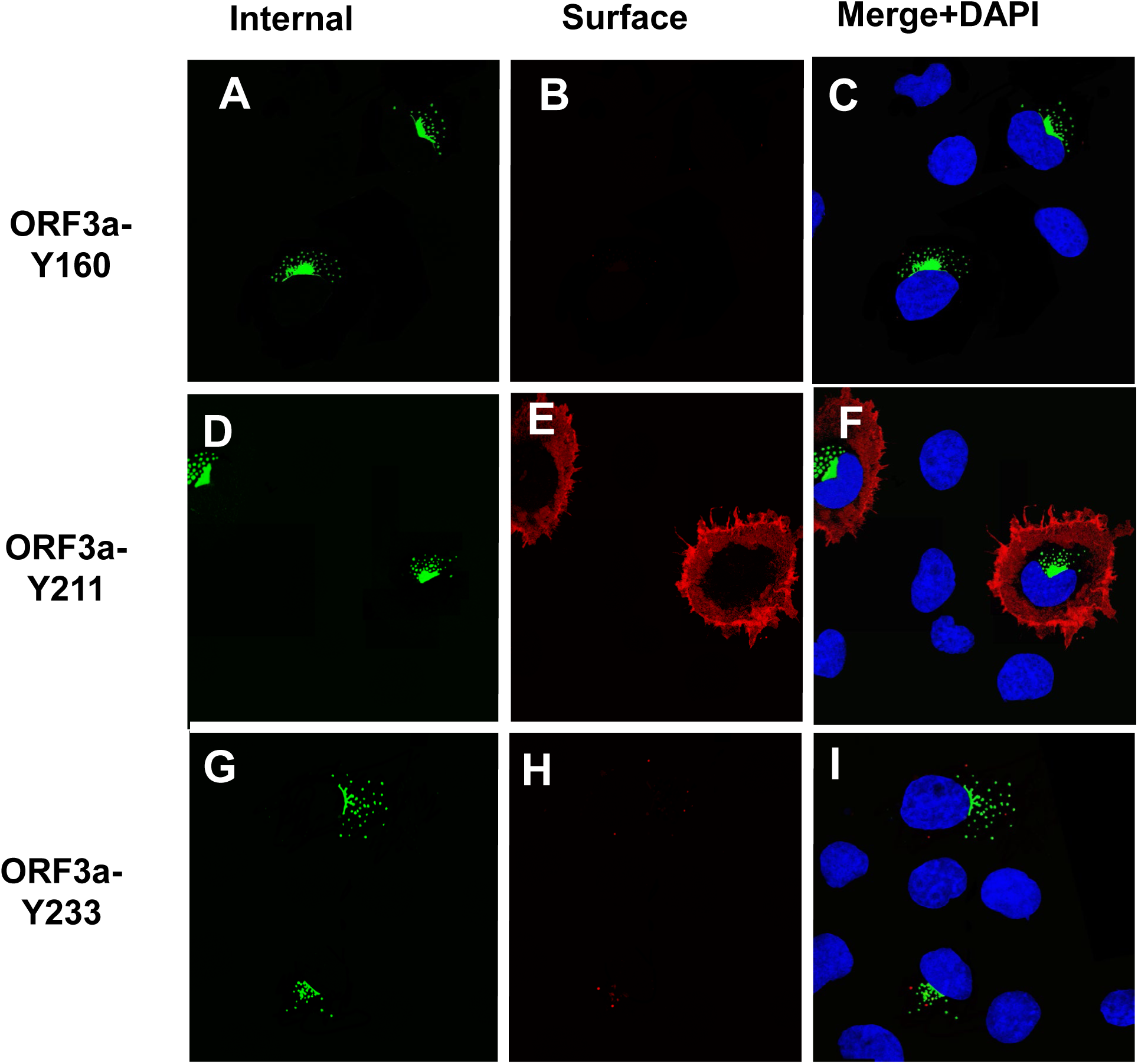
Surface immunostaining of cells transfected with vectors expressing ORF3a mutants with one tyrosine motif intact. COS-7 cells were transfected with vectors expressing each of the ORF3a mutants. At 24 h post-transfection, cells were immunostained with an antibody against the HA-tag followed by a secondary antibody tagged with Alexa Fluor 594. The cells were washed three times, and permeabilized as described in the Materials and Methods section. Permeabilized cells were then reacted with the same primary antibody and a secondary antibody tagged with Alexa Fluor 488. The cells were washed, mounted, and examined using a Leica TSP8 laser scanning confocal microscope. Shown are individual red and green images and merged images of the red, green, and blue channels. Panel A. Surface immunostaining of HA-ORF3a-Y160 with an antibody against the HA-tag. Panel B. Internal immunostaining of HA-ORF3aY160 with an antibody against the HA-tag. Panel C. Merge of Panels A and B. Panel D. Surface immunostaining of HA-ORF3a-Y211 with an antibody against the HA-tag. Panel E. Internal immunostaining of HA-ORF3a-211 with an antibody against the HA-tag. Panel F. Merge of Panels D and E. Panel G. Surface immunostaining of HA-ORF3a-Y233 with an antibody against the HA-tag Panel H. Internal immunostaining of HA-ORF3a-233 with an antibody against the HA-tag Panel I. Merged of cells Panels G and H.

**Figure 9.**
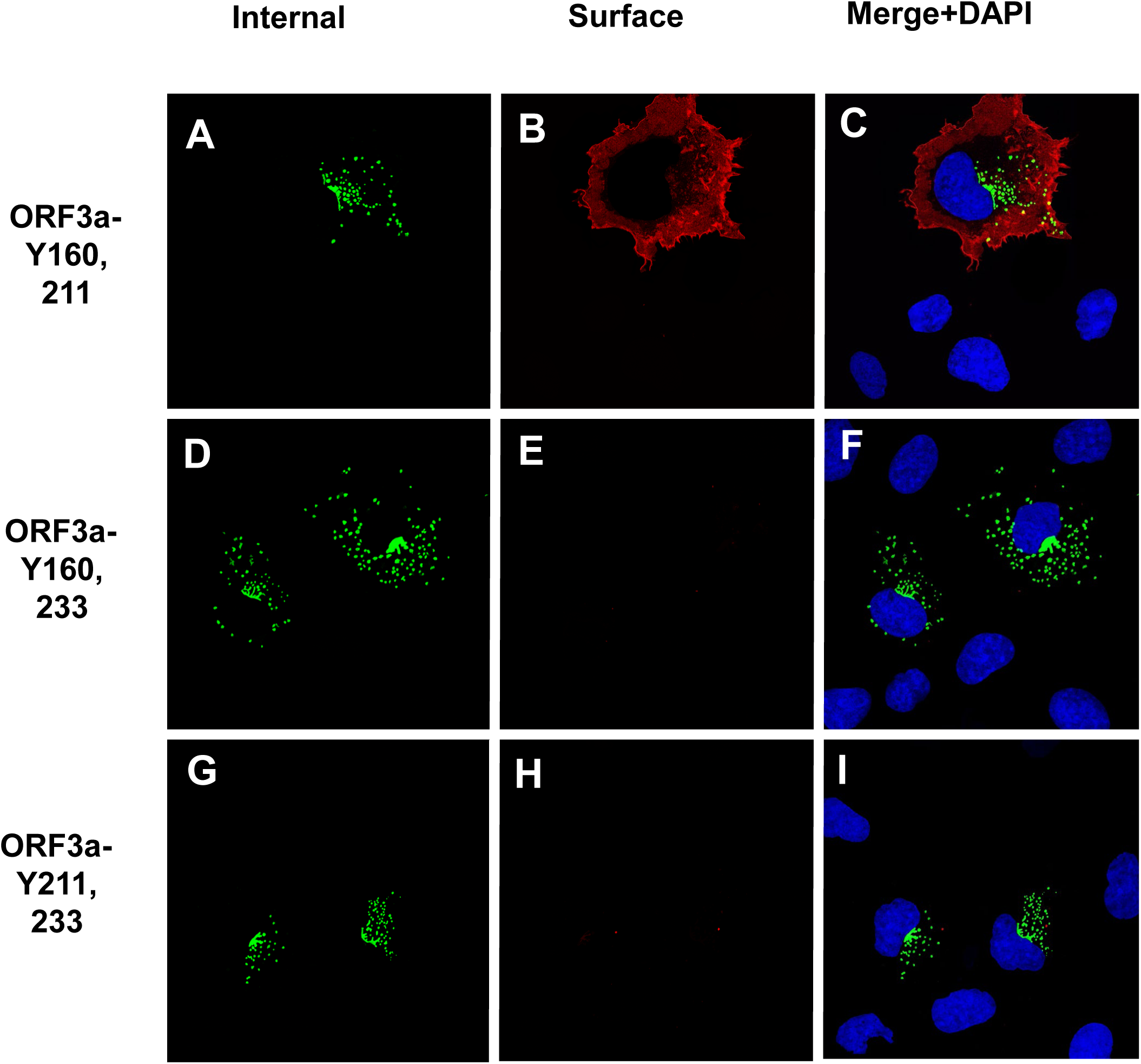
Surface immunostaining of cells transfected with vectors expressing ORF3a mutants with two tyrosine motifs intact. COS-7 cells were transfected with vectors expressing each of the ORF3a mutants. At 24 h post-transfection, cells were immunostained with an antibody against the HA-tag followed by a secondary antibody tagged with Alexa Fluor 594. The cells were washed three times and permeabilized as described in the Materials and Methods section. Permeabilized cells were then reacted with the same primary antibody and a secondary antibody tagged with Alexa Fluor 488. Cells were washed, mounted, and examined using a Leica TSP8 laser scanning confocal microscope. Shown are individual red and green images and merged images of the red, green, and blue channels. Panel A. Surface immunostaining of HA-ORF3a-Y160,211 with an antibody against the HA-tag. Panel B. Internal immunostaining of HA-ORF3a-Y160,211 with an antibody against the HA-tag. Panel C. Merge of Panels A and B. Panel D. Surface immunostaining of HA-ORF3a-Y160,233 with an antibody against the HA-tag. Panel E. Internal immunostaining of HA-ORF3a-Y160,233 with an antibody against the HA-tag. Panel F. Merge of Panels D and E. Panel G. Surface immunostaining of HA-ORF3a-Y211,233 with an antibody against the HA-tag Panel H. Internal immunostaining of HA-ORF3a-Y211,233 with an antibody against the HA-tag. Panel I. Merged of cells Panels G and H.

#### Substitution of the tyrosine residues in the three motifs with phenylalanine residues

A potential caveat to the above results is that substitution of the tyrosine residues with alanines may have altered the structure of the cytoplasmic domain resulting in the observed results. To address this potential concern, we substituted the tyrosine residues of the three potential tyrosine motifs with structurally similar phenylalanine residues. Previously, it was shown that tyrosine could not be effectively substituted by the structurally similar phenylalanine (**20**). These three constructs, designated as HA-ORF3a-Y160F, HA-ORF3a-Y211F, and HA-ORF3a-Y233F (with the other two tyrosine motifs intact) were analyzed for ORF3a expression in the RER, TGN, and at the cell plasma membrane. The results indicate that HA-ORF3a-Y160F, HA-ORF3a-Y211F, and HA-ORF3a-Y233F had a similar intracellular localization pattern as the HA-ORF3a-Y211,233, HA-ORF3a-Y160,233, and ORF3a-Y160,211, respectively (**Fig. 10**). These results argue that the single tyrosine to alanine substitutions likely did not alter the overall structure of the cytoplasmic domain.

**Figure 10.**
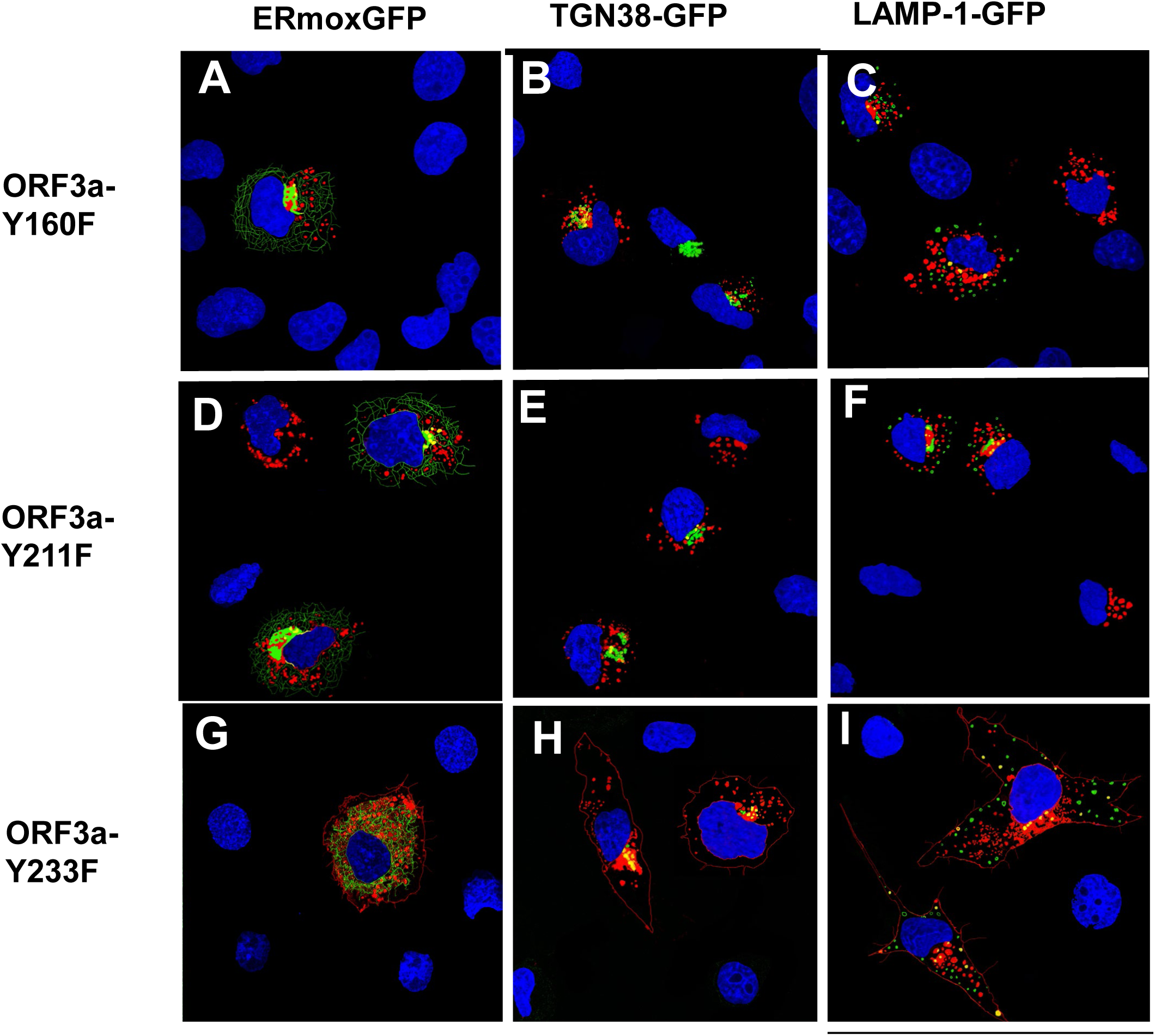
Phenylalanine residues cannot substitute for tyrosine residues in the tyrosine motifs. COS-7 cells were grown in 6-well plates with coverslips. Cells (70% confluent) were in were co-transfected with vectors expressing HA-ORF3a-Y160F Panels A-C), HA-ORF3a-Y211F (Panels D-F), or HA-ORF3a-Y233F (Panels G-I) and either ERmoxGFP (Panels A,D, G) or TGN-38GFP (Panels B,E,H) LAMP-1-GFP (panels C, F, I). At 48 h post-transfection, the cells were processed for immunofluorescence as described in Figure 3.

#### Lysosome localization of the ORF3a mutants

We next analyzed the unmodified ORF3a, and ORF3a mutants for co-expression with LAMP-1, a protein associated with late endosomes and lysosomes. Vectors expressing the unmodified ORF3a-HA or each mutant were transfected into COS-7 cells and at 48h post-transfection, cells were fixed and permeabilized followed by immunostaining for the HA-tag on the ORF3a protein and for LAMP-1. Laser scanning confocal microscopy was used to examine the cells for LAMP-1 and ORF3a proteins, respectively. The results of the confocal microscopy revealed that expression of the unmodified ORF3a co-localized with LAMP-1 in the region of the ER, and in vesicular structures closer to the cell plasma membrane (**Fig. 11**). In contrast, cells transfected with vectors expressing HA-ORF3a-ΔYxxΦ, HA-ORF3a-Y160, HA-ORF3a-Y233, HA-ORF3a-Y211,233 and HA-ORF3a-Y160,233 co-localized with LAMP-1 in the ER region of the cell but none of these mutants co-localized with LAMP-1 positive lysosomes towards the periphery of the cell (**Fig. 11**). Finally, HA-ORF3a-Y211 and HA-ORF3a-Y160,211 were like the unmodified HA-ORF3a in their pattern of co-localization with LAMP-1, suggesting that targeting of ORF3a to the lysosomes likely required the tyrosine motif ^211^YYQL^213^.

**Figure 11.**
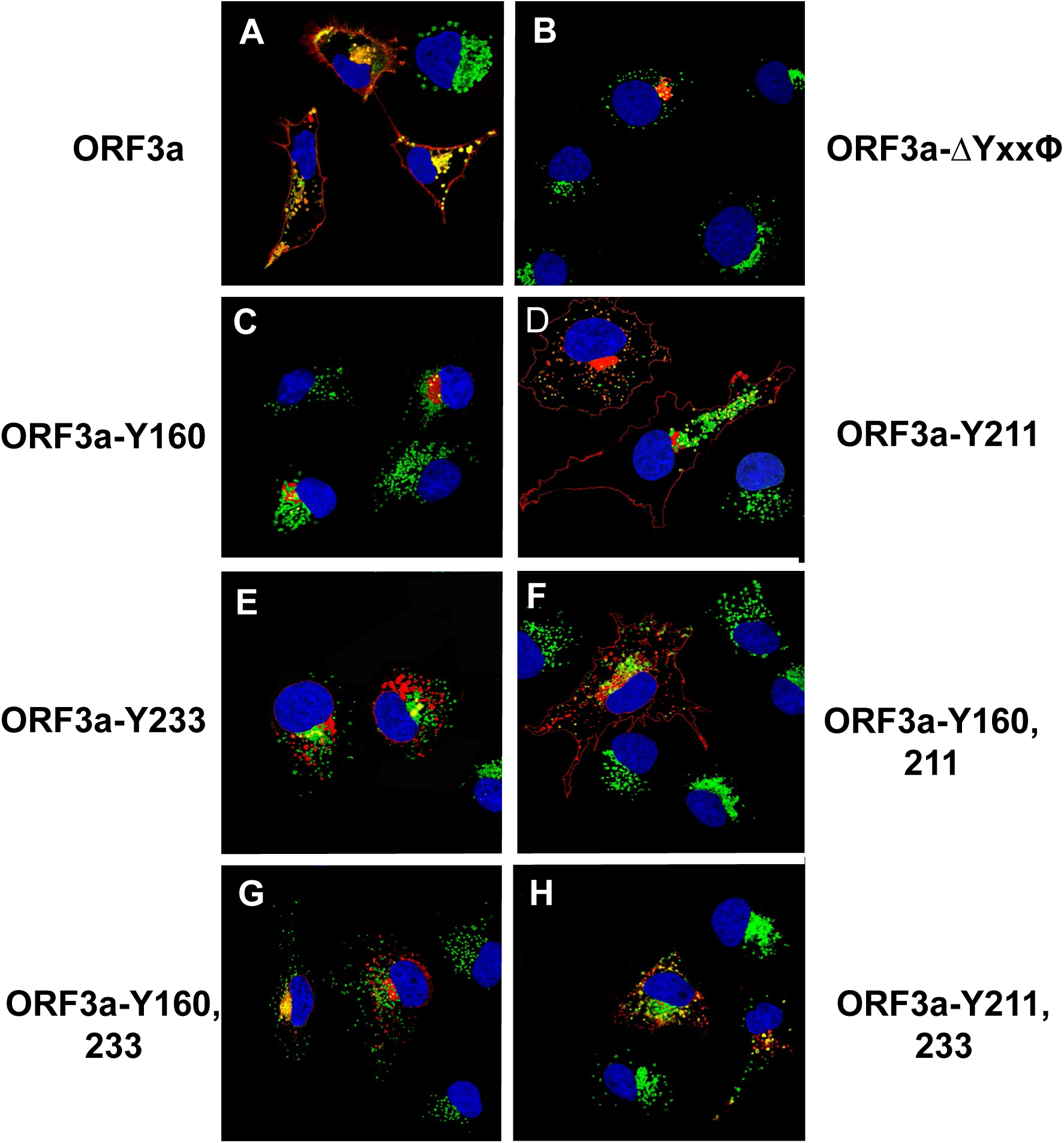
Co-localization of ORF3a and the tyrosine-based sorting signal mutants with LAMP-1. COS-7 cells were transfected with the empty vector or the same vector expressing ORF3a or individual ORF3a mutants. At 48 hr. post-transfection, the cells were fixed, permeabilized, and blocked. Cells were reacted with a mouse monoclonal antibody against the HA-tag and a rabbit antibody against LAMP-1 overnight, washed, and reacted with an appropriate secondary antibody tagged with Alexa Fluor 594 (for HA) and Alexa Fluor 488 (for LAMP-1) for 1 h. Cells were washed and counter-stained with DAPI (1 μg/ml) for 5 min. Cells were viewed using a Leica TC8 confocal microscope and at least 50 cells were examined for co-localization with the LAMP-1 marker. Panel A. HA-ORF3a. Panel B. HA-ORF3a-YxxΦ. Panel C. HA-ORF3a-Y160. Panel D. HA-ORF3a-Y211. Panel E. HA-ORF3a-Y233. Panel F. HA-ORF3a-Y160,211. Panel G. HA-ORF3a-Y160, 233. Panel H. HA-ORF3a-Y211,233.

### Induction of apoptosis by the ORF3a mutants

ORF3a was previously demonstrated to cause both intrinsic and extrinsic apoptosis (**6**). Further, these investigators observed that mutation of the tyrosine at position 160 to alanine (within the **<u>Y</u>**NSV motif) eliminated ORF3a-induced apoptosis. We examined whether ORF3a and the seven ORF3a mutants resulted in the cleavage of procaspase 3, which is an effector caspase of both the intrinsic and extrinsic pathways of apoptosis. HEK293 cells were transfected with the empty vector and used as a background control for procaspase 3 cleavage, while cells transfected with the empty vector and treated with 3 μM staurosporine (STS) for 18 hours served as a positive control for cleavage of procaspase 3 activity. The assays were run a total of four times. Our results indicated significant procaspase 3 cleavage activity in the cells transfected with the empty pcDNA3,1(+) vector and treated with 3 μM STS while cells transfected with the empty pcDNA3.1(+) showed levels like non-transfected cells (**Fig. 12**). Cells transfected with the vector expressing ORF3a had low levels of procaspase 3 cleavage although it was not statistically significant (*p* value of 0.54). Similarly, transfection of cells with vectors expressing the HA-ORF3a-ΔYxxΦ, HA-ORF3a-Y160, and HA-ORF3a-Y211 mutants had no procaspase 3 cleavage activity above that of the pcDNA3.1(+) control while HA-ORF3a-Y233, HA-ORF3a-Y160,233, HA-ORF3a-Y211,233, and HA-ORF3a-Y160,233 mutants had low levels of procaspase 3 cleavage activity (**Fig. 12**). These results suggest that the tyrosine motif ^233^YNKI^236^ may influence the level of procaspase 3 cleavage activity, although the level of activity was not considered to be statistically significant.

**Figure 12.**
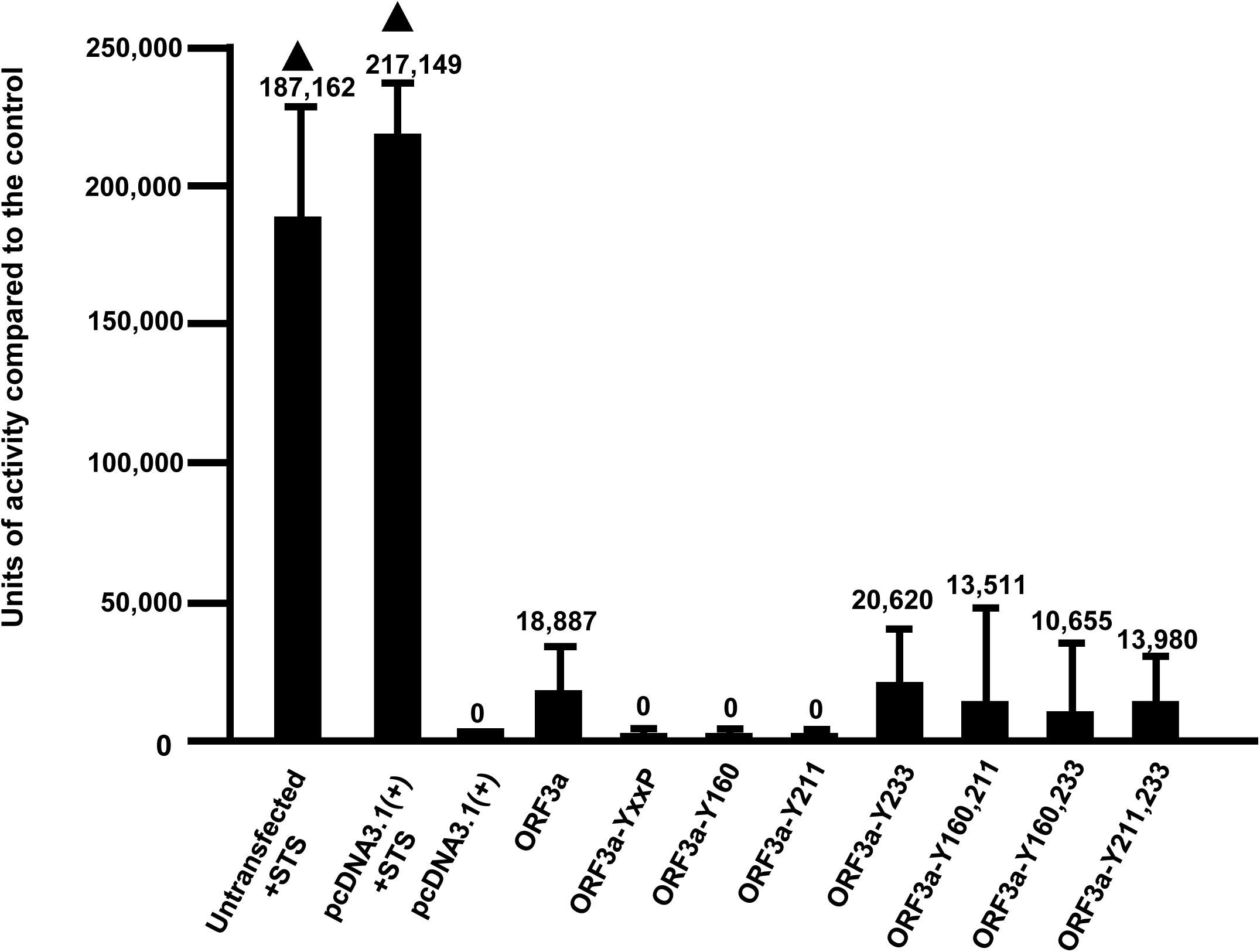
The role of the potential tyrosine-based sorting signals of ORF3a on the induction of apoptosis. HEK293 cells were either not transfected, transfected with empty vector pcDNA3.1, or the same vector expressing the unmodified ORF3a, or the tyrosine motif mutants. At 48 h, cells were assayed for the presence of caspase 3 activity using the EnzChekTM Caspase-3 Assay Kit #1 according to the manufacturer’s instructions. Controls included transfected cells treated with 2 μM staurosporine (STS) for 18h, and transfected cells treated with the pan-caspase inhibitor Z-VAD-FMK (InvivoGen). The assays were performed a minimum of three times and analyzed for statistical significance using Students’ *t*-test.

### The role of the ORF3a tyrosine motifs in the induction of autophagy

In previous studies, it was reported that ORF3a can induce incomplete autophagy (**7, 21, 22**). In the initial steps of autophagy, LC3B-I is lipidated with phosphatidylethanolamine (PE) resulting in the formation of LC3B-II, which is generally accepted as the gold standard for the initiation of autophagy. Adaptor protein SQSTM1/p62 binds to ubiquitinated proteins and LC3-II for mediating autophagy by localizing ubiquitinated proteins and organelles in autophagosomes. Interestingly, while ORF3a increases LC3-II formation, apparently there is no degradation of p62, which would be expected if autophagosomes did not fuse with lysosomes to form autophagolysosomes and cargo degraded. We examined the ORF3a mutants for LC3-II formation and levels of SQSTM1/p62. Compared with HEK293 cells transfected with the empty vector, transfection of cells with the vector expressing unmodified ORF3a resulted in an increase in LC3-II but with little degradation of p62. These results were comparable to previous studies (**7, 21, 23**). Analysis of ORF3a and the ORF3a mutants for LC3-II lipidation indicated that ORF3a and the tyrosine mutants had increased lipidation of the LC3-I to LC3-II when compared with the empty vector control (**Fig. 13**). Examination of cells transfected with vectors expressing the unmodified ORF3a or ORF3a with amino acids substitutions of the tyrosine residues revealed that p62 was not degraded in cells but was increased over the empty vector control (**Fig. 13**).

**Figure 13.**
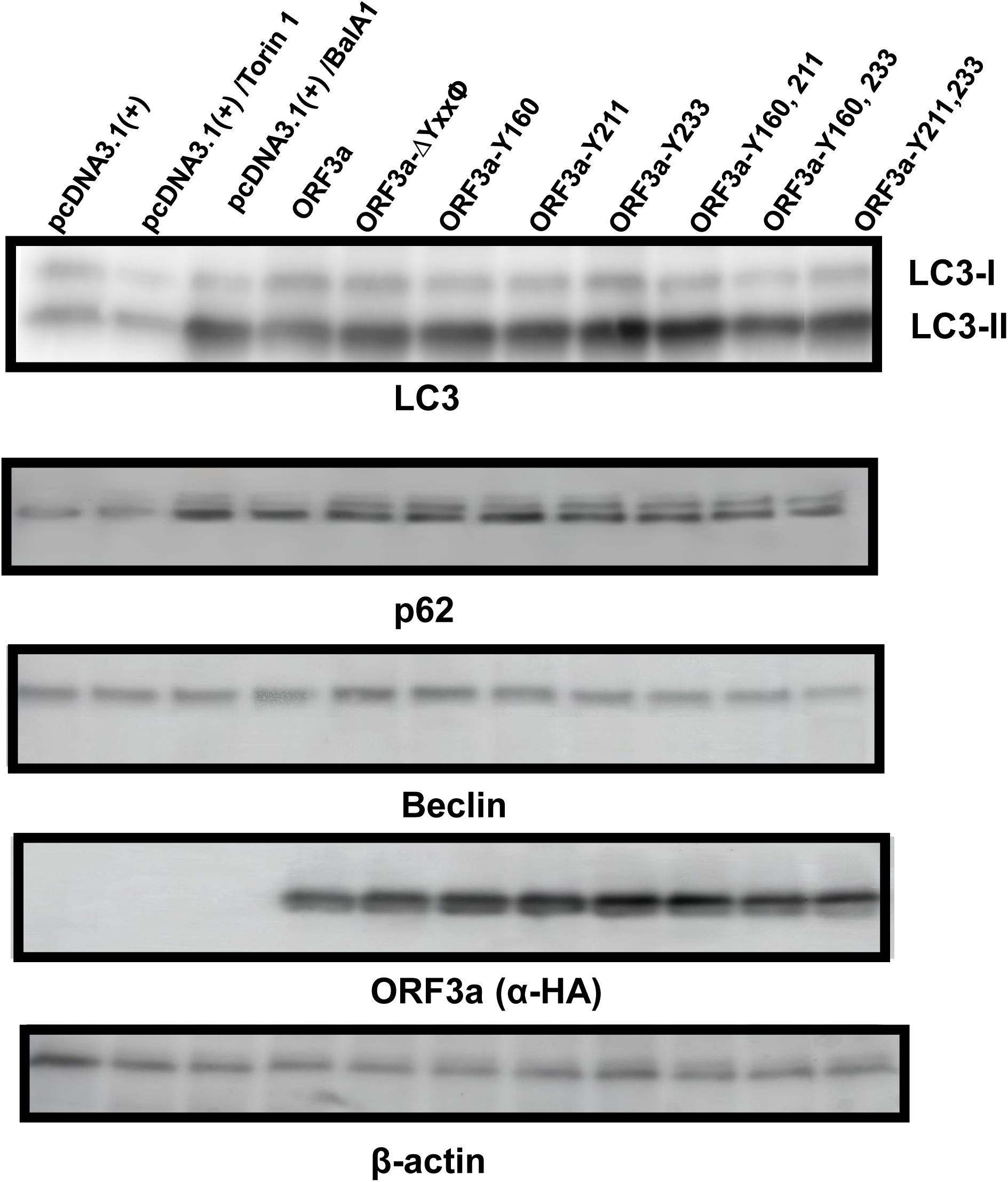
The role of the different tyrosine motifs in the induction of autophagy. HEK293 cells were transfected with vectors expressing the unmodified ORF3a or the six tyrosine mutants. Controls included transfection with empty pcDNA3.1(+) vector alone and transfection in the presence of Torin or balfinomycin A. At 48 h post-transfection, cells were washed, pelleted, and lysed in 2x sample-reducing buffer. The lysates were subjected to SDS-PAGE and proteins transferred to PVDF membranes as described in the Materials and Methods section and analyzed by immunoblots using antibodies to LC3, p62, Beclin, ORF3a, or β-actin. Panel A. Analysis of LC-I and LC-II. Panel B. Analysis of p62 levels. Panel C. Analysis of Beclin levels. Panel D. Analysis of ORF3a expression. Panel E. Analysis of β-actin expression (for loading control).

## DISCUSSION

Previous studies have shown that the ORF3a viroporin of SARS-CoV-2 is a virulence factor in pathogenesis (**4**). Using the hACE-2/mouse model, deletion of ORF3a from SARS-CoV-2 did not result in a significant reduction in virus titers in cell culture but had a profound effect on *in vivo* lung pathology (**4**), indicating that the ORF3a contributes to *in vivo* pathology. ORF3a has been reported to have several biological functions that could impact the pathogenesis in the hACE-2/mouse model. These functions include an ion channel activity, the ability to induce apoptosis, and to disrupt the autophagy pathway. Thus, identification of those protein domains that are important to ORF3a trafficking and biological functions *in vivo* could possibly aide in the development of more robust vaccines that result in long-term immunity against this viral disease.

In a recent study, the SARS-CoV was shown to cause apoptosis through both the intrinsic and extrinsic pathways while the SARS-CoV-2 ORF3a was shown to cause apoptosis through an extrinsic pathway (**6**). This was based on the findings that the presence of SARS-CoV-2 ORF3a resulted in the cleavage of pro-caspase 9 and pro-caspase 8 of the extrinsic pathway (**6**). These investigators showed that a tyrosine to alanine substitution within the tyrosine-based sorting motif (^160^YNSV^163^) resulted in a protein (ORF3a-YA) that no longer trafficked to the cell plasma membrane and did not induce apoptosis, suggesting that cell surface expression was necessary for SARS-CoV-2 ORF3a induced apoptosis. However, these investigators also showed that ORF3a-YA was not associated with membrane fractions of the cell. This is puzzling as this tyrosine to alanine substitution should not affect the biosynthesis and translocation of this protein across the RER membrane. Further, it is unclear why these investigators did not analyze the other potential tyrosine-based sorting motifs within the cytoplasmic domain.

In this study, we have expanded on the studies of the tyrosine-based motifs of the SARS-CoV-2 ORF3a protein. Analysis of the cytoplasmic domain (CD) of these ORF3a proteins revealed that more than one potential tyrosine-based sorting motif exists in the CD of these proteins (**SARS-CoV-2**: ^160^YNSV^163^, ^211^YYQL^214^, and ^233^YKNI^236^, **SARS-CoV**: ^160^YNSV^163^, ^200^YVVV^203^, and ^211^YYQL^214^). The central question we addressed is whether one or more of these other potential tyrosine motifs contribute to the trafficking of the SARS-CoV-2 ORF3a to the cell surface and lysosomes, and if so, do they also contribute to the biological functions of this protein? We used a strategy in which we first eliminated all three tyrosine-based sorting signals by altering each tyrosine to an alanine. The resulting protein, ORF3a-ΔYxxΦ, was detectable in several organelles of the secretory pathway (ER, ERGIC, *cis-medial* Golgi, *trans* Golgi and TGN). However, ORF3a-ΔYxxΦ was neither detectable at the cell surface nor did it co-localize with LAMP-1, a marker for late endosomes/lysosomes. Conversely, the intact ORF3a was easily detected at the cell plasma membrane. To determine the importance of each individual tyrosine motif alone or the combinations of two motifs, we generated mutants with one or two tyrosine-based sorting motifs intact. For analysis of intracellular trafficking, we used either: a) plasmids expressing proteins tagged with fluorescent proteins that localize in different organelles; or b) antibodies specific for organelle specific proteins. We reasoned that if the ^160^YNSV^163^ motif was critical to transport to the surface, the mutant with just ^160^YNSV^163^ motif intact (ORF3a-Y160) should be transported to the cell surface. Our results indicated that ORF3a-Y160 was inefficiently transported to the cell surface, indicating that the ^160^YNSV^163^ motif by itself does not dictate efficient transport to the cell surface. Our results indicated that the ORF3a with only the ^211^YYQL^214^ motif intact (ORF3a-Y211) was efficiently transported to the cell surface. Finally, our data also indicated that mutant ORF3a-Y160,211 was also transported to the cell surface suggesting that these two motifs may act cooperatively to enhance the transport of ORF3a to the cell surface or they do not interfere each other. Finally, the ^233^YKNI^236^ motif did not appear to be involved in transport to the cell plasma membrane. This is based on the results with ORF3a-Y160,233, ORF3a-211,233, and ORF3a-Y233, which were not detected at the cell plasma membrane. Also, the tandem presence of Y211 and Y233 interfered with the transport of ORF3a-Y211,233 to the plasma membrane. This suggests that the Y233 motif may interfere in the trafficking of ORF3a to the cell surface in the presence of one other tyrosine motif (211) and that the presence of both Y160 and Y211 motifs is required to overcome the influence of Y233 motif. Alternatively, the Y233 motif may be necessary for endocytosis, perhaps to the lysosomes of the cell, which has been well documented (**7, 21, 25**). Thus, while the results of Ren and colleagues (**6**) were correct with their Y160A mutant (the equivalent to our ORF3a-Y211,233 mutant, it was likely not due to the disruption of the ^160^YNSV^163^ motif. One caveat to our studies is that the substitution of the tyrosine residues with alanines may have changed the overall structure of the CD. However, comparable results were obtained when we substituted the tyrosine residues with structurally similar phenylalanines, suggesting that an overall change in the CD was not likely the cause for our results (**Fig. 10**). While our studies concentrated on the ORF3a of SARS-CoV-2, it is of interest that of the three tyrosine motifs examined in 14 SARS-CoV-2 and SARS-CoV-2-like isolates, the ^160^YNSV^163^ motif was 100% conserved while the ^211^YYQL^214^ motif was conserved in 13 of 14 isolates (∼92%) examined.

Autophagy is a conserved cellular process of intracellular degradation of senescent or malfunctioning organelles to maintain intracellular homeostasis (**26-29**). Autophagy occurs in response to different forms of stress, including nutrient deprivation, growth factor depletion, infection, and hypoxia. This process can target viral components or even full viruses for lysosomal degradation (**30**). Most successful viruses developed strategies to avoid degradation by autophagy or have evolved to exploit components of the autophagic machinery to enhance their replication and to mediate membrane trafficking and fusion processes. In a recent study, the role of different SARS-CoV-2 proteins revealed that E, M, ORF3a, ORF7a and Nsp15 affected autophagy (**31,32**). These investigators found that the number of LC3-II positive autophagosomes was decreased in the presence of Nsp15 while expression of E, M, ORF3a, and ORF7a caused a strong accumulation of membrane-associated LC3-II (**31**). ORF3a increases the conversion of microtubule-associated protein light chain 3 (LC3B-I) to the lipidated form, LC3B-II, while the level of SQSTM1/p62 did not decrease, indicating that ORF3a blocks autophagosome-lysosome fusion. Relating to this incomplete autophagy, late endosomal ORF3a interacts directly with and sequestrates VPS39 of the homotypic fusion and protein sorting (HOPS) complex (**7, 21**). This prevents the HOPS complex from interacting with the autophagosomal SNARE protein STX17 and blocks the assembly of the STX17-SNAP29-VAMP8 SNARE complex, which mediates autophagosome/ amphisome fusion with lysosomes (**22, 33**). Finally, one study has suggested that ORF3a may mediate virus release from lysosomes. However, the extent to which this occurs within infected cells is still unknown (**34**). We analyzed LC3-II lipidation of the unmodified ORF3a and our mutants. Our results indicated that lipidation of LC3-I was similar in ORF3a-transfected and ORF3a mutant transfected cells, indicating that the tyrosine motifs did not play a crucial role in this process. Upon fusion of LC3-II decorated autophagosomes with lysosomes, the autophagic receptor p62 is degraded (autophagic turnover). Analysis of p62 levels in cells expressing the unmodified ORF3a or the tyrosine motif mutants was essentially the same with the unmodified ORF3a and the tyrosine motif mutants, regardless of if they were transported to the cell surface or retained in intracellular compartments.

In addition to its role in transport to the cell plasma membrane, the same sorting signals are also involved in targeting membrane proteins to the lysosomes (**13, 35, 36**). Previous studies have shown that ORF3a interacts with the components of the lysosome to prevent fusion of lysosomes with autophagosomes (**7, 21, 25**). Lysosomal membrane proteins are targeted to lysosomes using either direct or indirect pathways. With the direct pathway, lysosomal membrane proteins are transported from the TGN to either early or late endosomes and then to lysosomes. In the indirect pathway, lysosomal proteins are first transported from the TGN to the plasma membrane followed by endocytosis to early endosomes and eventual delivery to late endosomes/lysosomes. Lysosomal membrane proteins possess sorting signals in their cytoplasmic domains that mediate both lysosomal targeting and rapid endocytosis from the cell surface. These signals have been best characterized for members of the lysosomal-associated membrane proteins (LAMP) and lysosomal integral membrane proteins (LIMP) but are also present in other lysosomal membrane proteins. Like other TGN sorting and endocytic signals, the majority of lysosomal targeting signals belong to either the YxxΦ or [DE]xxxL[LI] types but also have other features that make them functional for lysosomal targeting. One of the most important features is the placement of either type of signal close (often 6–13 residues) to the transmembrane domain (**37, 38**). The use of multiple YxxΦ signals for targeting proteins to the lysosomes has been reported. SID1 transmembrane family member 2 (SDIT2) is an integral membrane protein of lysosomes that mediates the translocation of RNA and DNA across the lysosomal membrane during RNA and DNA autophagy (RDA), a process in which RNA or DNA is directly imported into lysosomes and degraded (**39, 40**). Human and mouse SIDT2 homologs show 95% sequence identity across the entire protein (832 amino acids) and 100% identity at the C-terminal 100 amino acids (**41**). With SIDT2, localization to the lysosomal membrane was mediated by three cytosolic YxxΦ motifs located between transmembrane 1 and 2, and SIDT2 interacts with AP-1 and AP-2 through the Y359GSF motif (**42**). Our data indicated that tyrosine-based sorting signals (YxxΦ) at positions 160 and 211 were necessary for ORF3a trafficking to lysosomes, which coincidentally were the same motifs required for transport to the cell plasma membrane. This suggests that ORF3a is likely transported to the cell surface prior to endocytosis and targeting lysosomes. This is of importance as it was recently reported that β-coronaviruses such as SARS-CoV-2 use the endosomal pathway and lysosomes for egress rather than the secretory pathway (**34**). In this study, the investigators showed that ORF3a co-localized with lysosomes and presented evidence that ORF3a caused lysosome de-acidification, presumably through its viroporin activity. This was determined by the expression of ORF3a and staining transfected cells with Lysotracker Red DND-99, which is a cell permeable, acidophilic dye that accumulates in acidic organelles. Under these conditions, the Lysotracker red fluorescence was diminished in the presence of ORF3a, suggesting that ORF3a may have caused the de-acidification of the lysosomes via a proton channel. Whether ORF3a is a proton ion channel remains to be determined. Recently, SARS-CoV-2 ORF3a was expressed in *Spodoptera frugiperda*, reconstituted into liposomes, and single-channel currents were recorded from excised patches (**4**). ORF3a behaved as a cation channel with a large single-channel conductance (375 pA) that had a modest selectivity for Ca^+2^ and K^+^ over Na^+^ ions and was not blocked by Ba^++^, which was the case for the SARS-CoV channel (**26**). However, no studies have reported SARS-CoV-2 ORF3a being a proton ion channel. This differed from the SARS-CoV ORF3a, which could induce apoptosis via the intrinsic pathway. The apoptosis function of SARS-CoV ORF3a has been reported to involve the ion channel activity of this protein (**43**).

In addition to potential effects of the ORF3a tyrosine motifs on the intracellular transport to the cell surface and lysosomes, the SARS-CoV ORF3a induces NF-kB activation, chemokine production, Golgi fragmentation, accumulation of intracellular vesicles, and cell death (**44, 45**). Both SARS-CoV and SARS-CoV-2 ORF3a proteins have been implicated in the induction of apoptosis (**6, 44, 46, 47**). In the most recent study, investigators found a correlation between SARS-CoV-2 ORF3a-induced apoptosis and cell plasma membrane expression (**6**). Mutation of the tyrosine of ^160^YNSV^163^ resulted in neither plasma membrane expression nor apoptosis (**6**). However, the use of established cellular markers to determine the intracellular site of expression was not performed. We analyzed the level of apoptosis caused by the unmodified ORF3a and various tyrosine mutants. We analyzed the levels of cleavage of procaspase 3 to caspase 3, which is an effector caspase. We observed that ORF3a-Y160, which we found was not observed at the cell surface, induced caspase 3 activity while mutant ORF3a-Y211, which was expressed at the cell surface, did not induce caspase 3 activity, indicating there was no correlation between cell surface expression and apoptosis. This was reinforced with other ORF3a mutants (**Fig. 12**).

As discussed earlier, deletion of the ORF3a gene results in a SARS-CoV-2 that is less pathogenic in the K18 mouse model of SARS-CoV-2 pathogenesis (**48**). However, the role of individual motifs in the biological functions of ORF3a in pathogenesis such as the tyrosine-based sorting signals examined here have yet to be addressed. Elucidation of such motifs in ORF3a and their role in pathogenesis along with the identification of critical motifs in other genes of SAR-CoV-2 may lead to the development of live attenuated vaccines that lead to better and longer-term immunity than current vaccines.

## MATERIALS AND METHODS

### Cells, viruses and plasmids

HEK293 and COS-7 cells were used for transfection of vectors expressing coronavirus proteins. Both cell lines were maintained in Dulbecco’s minimal essential medium (DMEM) with 10% fetal bovine serum (R10FBS), 10 mM Hepes buffer, pH 7.3, 100 U/ml penicillin, 100 μg/ml streptomycin and 5 μg gentamicin. Plasmids (all pcDNA3.1(+) based) expressing the SARS-CoV-2 ORF3a protein were synthesized by Synbio Technologies with a HA-tag at either a N-terminus (HA-ORF3a) or at the C-terminus (ORF3a-HA). Plasmids were sequenced to ensure that no deletions or other mutations were introduced during the synthesis. Expression of the ORF3a proteins was confirmed by transfection with the Turbofect transfection reagent (ThermoFisher) into 293 cells for 48 h followed by lysis of cells in 1X RIPA and immunoblot analysis using a mouse monoclonal antibody directed against the HA-tag (Thermo-Fisher, catalog # 26183; antibody 2-2.2.14). Other plasmids that expressed organelle markers tagged with fluorescent proteins were used for the intracellular localization of ORF3a proteins. These included: 1) ER-moxGFP for the rough endoplasmic reticulum (RER), Addgene catalog #68072 (a gift from Eric Snapp; 2) mNeonGreen-Giantin for cis-medial Golgi; Addgene catalog #98880 (a gift from Dorus Gadella); 3) TGN38-EGFP for *trans* Golgi network; Addgene catalog #128148 (a gift from Jennifer Lippincott-Schwartz); 4) 4xmts-mNeonGreen for mitochondria; Addgene catalog #98876 (a gift from Dorus Gadella); and 5LAMP-1-mNeonGreen for lysosomes; Addgene #98882 (a gift from Dorus Gadella).

### Site-directed mutagenesis of ORF3a

For site-directed mutagenesis, the pcDNA3.1(+) vector containing the SARS-CoV-2 HA-ORF3a gene was used in site-directed mutagenesis using a QuikChange II site-directed mutagenesis kit (Agilent) according to the manufacturer’s protocol. A similar mutant was constructed using ORF3a with C-terminal HA tag. We found no differences in the intracellular localization of the ORF3a with C- or N-terminal HA tags (data not shown). For construction of the ORF3a-ΔYxxΦ, the tyrosine residues of the three potential tyrosine signals (^160^YNSV^163^, ^211^YYQL^214^, and ^233^YNKI^236^) were changed to alanine residues (^160^**A**NSV^163^, ^211^**A**YQL^214^, ^233^**A**NKI^236^). The ORF3a-ΔYxxΦ gene was sequenced to ensure that the desired changes were made and that no unwanted changes occurred during the mutagenesis process. Using ORF3a-ΔYxxΦ, individual alanine residues were changed back to tyrosine residues to yield ORF3a-Y160, ORF3a-Y211, and ORF3a-Y233. ORF3a proteins with combinations of two tyrosine motifs were generated from the above single mutants to yield ORF3a-Y160,211, ORF3a-Y160,233, and ORF3a-Y211,233. Again, all were sequenced to ensure that the desired changes were made and that no unwanted changes occurred during the mutagenesis process.

### Immunofluorescence studies

To examine the intracellular localization of the SARS-CoV-2 ORF3a proteins, COS-7 cells grown on 13 mm coverslips were transfected with either the empty pcDNA3.1(+) vector, pcDNA3.1(+) expressing the unmodified ORF3a-HA, or the same vector expressing the various SARS-CoV-2 ORF3a mutants. Vectors were transfected into COS-7 cells using the Turbofect transfection reagent (ThermoFisher) according to the manufacturer’s instructions. With other experiments, vectors expressing unmodified ORF3a-HA or ORF3a mutants were co-transfected with vectors expressing various fluorescent marker proteins as described above. At 48 h post transfection, cells were washed three times in PBS, fixed in 4% paraformaldehyde (prepared in PBS) for 15 minutes, permeabilized with 0.1% Triton X-100 in PBS, and blocked for one hour with 22.5 mg/mL glycine and 0.1% BSA in PBST. The cultures were then incubated at 4C overnight with a mouse monoclonal antibody against HA-tag (Thermo-Fisher, antibody 2-2.2.14, #26183) and one of the following rabbit polyclonal or monoclonal antibodies: a) ERGIC53; b) Golgin-97 (trans Golgi marker; Abcam, ab84340) or c) LAMP-1 (late endosome/lysosome marker; CST, D2D11). The cells were washed in PBS and incubated with a secondary goat anti-rabbit antibody conjugated to AlexaFluor™-488 (Invitrogen, A11008) and a chicken anti-mouse conjugated to AlexaFluor™-594 (Invitrogen, A21201) for 1 h. Cells were counterstained with DAPI, and the coverslips were mounted on glass slides with ProLong^TM^ Diamond Antifade Mountant (ThermoFisher, P36961). The coverslips were viewed with a Leica TCS SP8 Confocal Microscope with a 100X objective and a 2X digital zoom using the Leica Application Suite X (LASX). A 405nm filter was used to visualize DAPI staining, a 488nm filter was used to visualize the organelle markers (to ER, ERGIC, cis/medial Golgi, trans Golgi, TGN, and late endosomes/lysosomes) and a 594nm filter was used to visualize the ORF3a-HA protein. To examine co-localization ORF3a proteins with mitochondria, COS-7 cells were co-transfected with vectors the ORF3a proteins and the vector or 4xmts-Neon-Green (mitochondria; Addgene, #98876). At 48 h post-transfection, cells were fixed, permeabilized and stained for ORF3a-HA proteins as described above. A minimum of 50 cells were examined for each sample, and the results presented in the figures are representative of each construct.

### Surface/internal immunofluorescence assays

To confirm the immunofluorescence results, we performed surface labeling experiments. COS-7 cells (2.5 x 10^5^) were plated onto a cover slip in 35 mm dishes overnight. Cells were washed and transfected with 1.5 ug of plasmid expressing the HA-ORF3a or mutants. At 24 h, cells were fixed in 4% freshly prepared paraformaldehyde for 15 min and washed three times with PBS, pH 7.4. The fixed cells were blocked with PBS containing 22.5 mg/mL glycine and 1% BSA for 1h, washed and incubated with the primary antibody (mouse anti-HA, 2-2.2.14 Invitrogen) in PBS containing 1% BSA at 1:400 dilution overnight at 4 C. Cells were washed three times and incubated with the first secondary antibody (chicken anti-mouse-AF594, A21201, Invitrogen) in PBS containing 1% BSA 1h at room temperature. Cells were permeabilized with PBS containing 0.2% Triton X-100 for 15 min and blocked with PBS containing 22.5 mg/mL Glycine, 1% BSA and 0.1% Tween-20. Cells were incubated with a 1:400 dilution of the primary antibody (mouse anti-HA, 2-2.2.14) in PBS containing 1% BSA and 0.1% Tween-20 for 1h at room temperature, washed three times and incubated with second secondary antibody (Rabbit anti-mouse-AF488, A11059, Invitrogen) in PBS containing 1% BSA 1h at room temperature. Finally, cells were washed three times, stained with DAPI 5 min and mounted using Prolong Diamond Antifade Mountant (Invitrogen, P36961). Cells were examined as described above.

### Apoptosis assays

We analyzed the ability of ORF3a-HA and the mutants for the ability to induce apoptosis in cells. We used a colorimetric caspase-3 assay protocol (EnzChek Caspase-3 Assay Kit #1, with the Z-DEVD-AMC substrate) based on the formation of the chromophore 7-amido-4-methyl coumarin (AMC) by cleavage from the labeled substrate Z-DEVD-AMC. HEK293 cells were transfected with either the empty vector, one expressing the unmodified ORF3a-HA or the ORF3a mutants. At 48 h, cells were lysed and assayed for caspase 3 activity according to the manufacturer’s instructions. The fluorescence of the AMC was quantified using a microtiter plate reader at 340 nm excitation and 441 nm emission.

## ACKNOWLEDGMENTS

We thank the ACGT for the sequence analyses and the KUMC Biotechnology Core for oligonucleotide synthesis. We also thank SynBio Technologies for the plasmid with the ORF3a gene. This work was supported by NIH R21AI158229 to ES/MK and by the KUMC Frontiers Clinical and Translational Science Awards Program 5UL1TR002366.

---

**Supplemental Figure 1.**
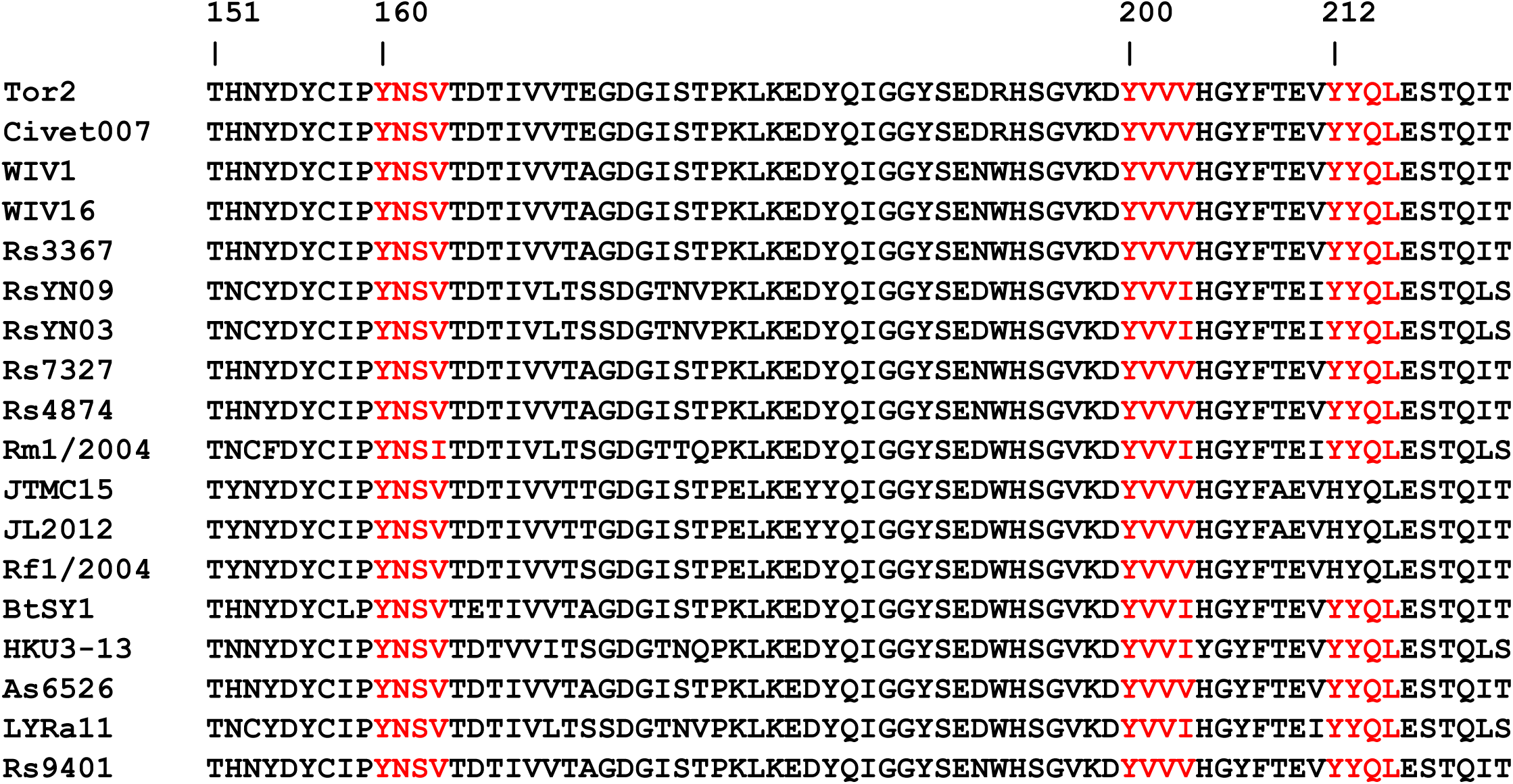
Eightteen ORF3a sequences from SARS-CoV (strain Tor2), Civet (Civet007) and SARS-CoV-like strains (all 274 amino acids in length). Shown are amino acids 160 to 240 with the potential tyrosine-based sorting motifs (in red). The species from which the isolate was obtained and accession numbers are: Tor2 (*Homo sapiens*; YP_009825052); Civet007 (*Paradoxurus hermaphroditus*; AAU04635); WIV1 (*Rhinolophus sinicus*; AGZ48832); WIV16 (*Rhinolophus sinicus*; ALK02458); Rs3367 (*Rhinolophus sinicus*; AGZ48819); RsYN09 (Rhinolophus stheno; QWN56264); RsYN03 (Rhinolophus sinicus; QWN56233); Rs7327 (*Rhinolophus sinicus;* ATO98219); Rs4874 (*Rhinolophus* sinicus; ATO98206); Rm1/2004 (*Rhinolophus macrotis*; ABD75326); JTMC15 (Rhinolophus ferrumequinum; ANA96028); JL2012 (*Rhinolophus ferrumequinum*;AIA62278); Rfl/2004 (*Rhinolophus ferrumequinum*; ABD75316); BtSY1 (*Rhinolophus thomasi;* WBV74274); HKU3-13 (*Rhinolophus spp*. ADE34824); As6526 (*Aselliscus stoliczkanus;* ATO98109); LYRa11 (Rhinolophus affinis; AHX37559); Rs9401 (*Rhinolophus sinicus;* ATO98232).

## REFERENCES

1. Tan YJ, Teng E, Shen S, Tan TH, Goh PY, Fielding BC, Ooi EE, Tan HC, Lim SG, Hong W. (2004). A novel severe acute respiratory syndrome coronavirus protein, U274, is transported to the cell surface and undergoes endocytosis. J Virol 78:6723–6734.

2. Lu W, Zheng BJ, Xu K, Schwarz W, Du LY, Wong CKL, Chen JD, Duan SM, Deubel V, Sun B. 2006. Severe acute respiratory syndrome-associated coronavirus 3a protein forms an ion channel and modulates virus release. Proc Natl Acad Sci 103:12540– 12545.

3. Chien TH, Chiang YL, Chen CP, Henklein P, Hänel K, Hwang IS, Willbold D, Fischer WB. 2013. Assembling an ion channel: ORF 3a from SARS-CoV. Biopolymers 99:628–635.

4. Kern DM, Sorum B, Mali SS, Hoel CM, Sridharan S, Remis JP, Toso DB, Kotecha A, Bautista DM, Brohawn SG. 2021. Cryo-EM structure of SARS-CoV-2 ORF3a in lipid nanodiscs. Nat Struct Mol Biol 28:573–582.

5. Miller AN, Houlihan PR, Matamala E, Cabezas-Bratesco D, Lee GY, Cristofori-Armstrong B, Dilan TL, Sanchez-Martinez S, Matthies D, Yan R, Yu Z, Ren D, Brauchi SE, Clapham DE. 2023.The SARS-CoV-2 accessory protein Orf3a is not an ion channel but does interact with trafficking proteins. Elife. 12:e84477.

6. Ren Y, Shu T, Wu D, Mu J, Wang C, Huang M, Han Y, Zhang XY, Zhou W, Qiu Y, Zhou X. 2020.The ORF3a protein of SARS-CoV-2 induces apoptosis in cells. Cell Mol Immunol 17:881–883.

7. Zhang Y, Sun H, Pei R, Mao B, Zhao Z, Li H, Lin Y, Lu K. 2021. The SARS-CoV-2 protein ORF3a inhibits fusion of autophagosomes with lysosomes. Cell Discov 7:31.

8. Anitei M, Hoflack B. 2011. Exit from the trans-Golgi network: from molecules to mechanisms. Curr Opin Cell Biol 23:443–451.

9. Bonifacino JS, Traub LM. 2003. Signals for sorting of transmembrane proteins to endosomes and lysosomes. Annu Rev Biochem 72:395–447.

10. Traub LM. 2003. Sorting it out: AP-2 and alternate clathrin adaptors in endocytic cargo selection. J Cell Biol. 163,203–208.

11. Kirchhausen T, Pines J, Toldo L, Lafont F. 1997. Membranes and sorting. Membrane permeability. Curr Opin Cell Biol. 9:473.

7. Kirchhausen T. 1999. Adaptors for clathrin-mediated traffic. Annu Rev Cell Dev Biol 15: 705–732.

8. Marks MS, Roche PA, Donselaar Evan, Woodruff L, Peters PJ, Bonifacino JS. 1995. A lysosomal targeting signal in the cytoplasmic tail of the beta chain directs HLA-DM to MHC class II compartments. J Cell Biol 131:351–369.

9. Marks MS, Ohno H, Kirchausen T, Bonifacino JS. 1997. Protein sorting by tyrosine based signals: adapting to the Ys and wherefores. Trends Cell Biol 7:124–128.

10. Rous BA, Reaves BJ, Ihrke G, Briggs JA, Gray SR, Stephens DJ, Banting G, Luzio JP. 2002. Role of adaptor complex AP-3 in targeting wild-type and mutated CD63 to lysosomes. Mol Biol Cell 13:1071–1082.

11. Stolt PC, Bock HH. 2006. Modulation of lipoprotein receptor functions by intracellular adaptor proteins. Cell Signal 18:1560–1571.

12. Traub, LM, Bonificano JS. 2013. Cargo recognition in clathrin-mediated endocytosis. Cold Spr Harb Perspect Biol 5: a016790.

18. Williams MA, Fukuda M. 1990. Accumulation of membrane glycoproteins in lysosomes requires a tyrosine residue at a particular position in the cytoplasmic tail. J Cell Biol 111:955–966.

19. Minakshi, R., Padhan, K. 2014. The YXXΦ motif within the severe acute respiratory syndrome coronavirus (SARS-CoV) 3a protein is crucial for its intracellular transport.

20. Boll W, Ohno H, Songyang Z, Rapoport I, Cantley LC, Bonifacino JS, Kirchhausen T.1996. Sequence requirements for the recognition of tyrosine-based endocytic signals by clathrin AP-2 complexes. EMBO J 15:5789–5795.

21. Miao G, Zhao H, Li Y, Ji M, Chen Y, Shi Y, Bi Y, Wang P, Zhang H. 2021. ORF3a of the COVID-19 virus SARS-CoV-2 blocks HOPS complex-mediated assembly of the SNARE complex required for autolysosome formation. Dev Cell 56:427–442.

22. Su WQ, Yu XJ, Zhou CM. 2021. SARS-CoV-2 ORF3a induces incomplete autophagy via the unfolded protein response. Viruses 13:2467.

23. Koepke L, Hirschenberger M, Hayn M, Kirchhoff F, Sparrer KM. 2021. Manipulation of autophagy by SARS-CoV-2 proteins. Autophagy 17: 2659–2661.

24. Vasquez DM, Park JG, Chiem K, Allué-Guardia A, Garcia-Vilanova A, Platt RN, Miorin L, Kehrer T, Cupic A, Gonzalez-Reiche AS, Bakel HV, García-Sastre A, Anderson T, Torrelles JB, Ye, Martinez-Sobrido L. 2021. Contribution of SARS-CoV-2 accessory proteins to viral pathogenicity in K18 human ACE2 transgenic mice. J Virol 95: e0040221.

25. Chen WJ, Goldstein JL, Brown MS. 1990. NPXY, a sequence often found in cytoplasmic tails, is required for coated pit-mediated internalization of the low density lipoprotein receptor. J Biol Chem 265:3116–3123.

26. Dikic I, Elazar Z. 2018. Mechanism and medical implications of mammalian autophagy. Nat Rev Mol Cell Biol 19:349–364.

27. Matsuzawa-Ishimoto Y, Hwang S, Cadwell K. 2018. Autophagy and Inflammation. Annu Rev. Immunol 36:73–101.

28. Mizushima N, Levine B. 2020. Autophagy in Human Diseases. N Engl J Med 383:1564–1576.

29. Shibutani ST, Saitoh T, Nowag H, Munz C, Yoshimori T. 2015. Autophagy and autophagy-related proteins in the immune system. Nat. Immunol 1:,1014–1024.

30. Choi, Y, Bowman JW, Jung, JU. 2018. Autophagy during viral infection — a double edged sword. Nature Rev Microbiol 16: 341–354.

31. Hahn M, Hirschenberger M, Koepke L, et al. 2021. Systematic functional analysis of SARS-CoV-2 proteins uncovers viral innate immune antagonists and remaining vulnerabilities. Cell Rep 35:109126.

32. Waisner H, Grieshaber B, Saud R, Henke W, Stephens EB, Kalamvoki M. 2023. SARS-CoV-2 harnesses host translational shutoff and autophagy to optimize virus yields: the role of the envelope (E) protein. Microbiol Spectr 11: e0370722.

33. Qu Y, Wang X, Zhu Y, Wang W, Wang Y, Hu G, Liu C, Li J, Ren S, Xiao MZX, Liu Z, Wang C, Fu J, Zhang Y, Li P, Zhang R, Liang Q. 2021. ORF3a-mediated incomplete autophagy facilitates severe acute respiratory syndrome coronavirus-2 replication. Front Cell Dev Biol 9:716208.

34. Ghosh S, Dellibovi-Ragheb TA, Kerviel A, Pak E, Qiu Q, Fisher M, Takvorian PM, Bleck C, Hsu VW, Fehr AR, Perlman S, Achar SR, Straus MR, Whittaker GR, de Haan CAM, Kehrl J, Altan-Bonnet G, Altan-Bonnet N. 2020. β-Coronaviruses use lysosomes for egress instead of the biosynthetic secretory pathway. Cell 183:1520–1535.

35. Braulke T, Bonifacino JS. 2009. Sorting of lysosomal proteins. Biochim Biophys Acta. 1793:605–614.

36. Höning S, J. Griffith J, H. Geuze H, W. Hunziker W. 1996. The tyrosine-based lysosomal targeting signal in LAMP-1 mediates sorting into Golgi-derived clathrin coated vesicles. EMBO J 15:5230–5239.

38. Rohrer J, Schweizer A, Russell D, Kornfeld S. 1996. The targeting of Lamp1 to lysosomes is dependent on the spacing of its cytoplasmic tail tyrosine sorting motif relative to the membrane. J Cell Biol 132:565–576.

39. Geisler C, Dietrich J, Nielsen B, Kastrup J, Lauritsen J, Odum N, Christensen M. 1998. Leucine-based receptor sorting motifs are dependent on the spacing relative to the plasma membrane. J Biol Chem 273: 21316–21323.

40. Jialin G, Xuefan G, Huiwen Z. 2010. SID1 transmembrane family, member 2 (Sidt2): a novel lysosomal membrane protein. Biochem Biophys Res Commun 402:588–594.

41 Aizawa S, Contu VR, Fujiwara Y, Hase K, Kikuchi H, Kabuta C, Wada K, Kabuta T. 2017. Lysosomal membrane protein SIDT2 mediates the direct uptake of DNA by lysosomes. Autophagy 13:218–222.

42. Nguyen TA, Smith BRC, Tate MD, Belz GT, Barrios MH, Elgass KD, Weisman AS, Baker PJ, Preston SP, Whitehead L, Garnham A, Lundie RJ, Smyth GK, Pellegrini M, O’Keeffe M, Wicks IP, Masters SL, Hunter CP, Pang KC. 2017. SIDT2 transports extracellular dsRNA into the cytoplasm for innate immune recognition. Immunity 47:498–509.

43. Contu VR, Hase K, Kozuka-Hata H, Oyama M, Fujiwara Y, Kabuta C, Takahashi M, Hakuno F, Takahashi SI, Wada K, Kabuta T. 2017. Lysosomal targeting of SIDT2 via multiple Yxxphi motifs is required for SIDT2 function in the process of Rnautophagy. J Cell Sci 130:2843–2853.

44. Siu KL, Yuen KS, Castaño-Rodriguez C, Ye ZW, Yeung ML, Fung SY, Yuan S, Chan CP, Yuen KY, Enjuanes L, Jin DY. 2019. Severe acute respiratory syndrome coronavirus ORF3a protein activates the NLRP3 inflammasome by promoting TRAF3-dependent ubiquitination of ASC. FASEB J 33:8865–8877.

45. Freundt EC, Yu L, Goldsmith CS, Welsh S, Cheng A, Yount B, Liu W, Frieman MB, Buchholz UJ, Screaton GR, Lippincott-Schwartz J, Zaki SR, Xu XN, Baric RS, Subbarao K, Lenardo MJ. 2010. The open reading frame 3a protein of severe acute respiratory syndrome-associated coronavirus promotes membrane rearrangement and cell death. J Virol 84:1097–109.

46. Kanzawa N, Nishigaki K, Hayashi T, Ishii Y, Furukawa S, Niiro A, Yasui F, Kohara M, Morita K, Matsushima K, Le MQ, Masuda T, Kannagi M. 2006. Augmentation of chemokine production by severe acute respiratory syndrome coronavirus 3a/X1 and 7a/X4 proteins through NF-kappaB activation. FEBS Lett 580:6807–6812.

47. Chan CM, Tsoi H, Chan WM, Zhai S, Wong CO, Yao X, Chan WY, Tsui SK, Chan HY. 2009. The ion channel activity of the SARS-coronavirus 3a protein is linked to its pro-apoptotic function. Int J Biochem Cell Biol 41:2232–2239.

48. Law PT, Wong CH, Au TC, Chuck CP, Kong SK, Chan PK, To KF, Lo AW, Chan JY, Suen YK, Chan HY, Fung KP, Waye MM, Sung JJ, Lo YM, Tsui SK. 2005. The 3a protein of severe acute respiratory syndrome-associated coronavirus induces apoptosis in Vero E6 cells. J Gen Virol 86:1921–1930.

49. Silvas JA, Vasquez DM, Park JG, Chiem K, Allué-Guardia A, Garcia-Vilanova A, Platt RN, Miorin L, Kehrer T, Cupic A, Gonzalez-Reiche AS, Bakel HV, García-Sastre A, Anderson T, Torrelles JB, Ye C, Martinez-Sobrido L. 2021. Contribution of SARS-CoV-2 accessory proteins to viral pathogenicity in K18 human ACE2 transgenic mice. J Virol 95: e0040221.

